# Sequencing of Monkeypox virus from infected patients reveals viral genomes with APOBEC3-like editing, gene inactivation, and bacterial agents of skin superinfection

**DOI:** 10.1101/2022.09.26.509493

**Authors:** Philippe Colson, Gwilherm Penant, Jeremy Delerce, Céline Boschi, Nathalie Wurtz, Stéphanie Branger, Jean-Christophe Lagier, Nadim Cassir, Hervé Tissot-Dupont, Matthieu Million, Sarah Aherfi, Bernard LA Scola

## Abstract

An epidemic of Monkeypox virus (MPX virus) infections has arisen in May 2022 in non-endemic countries particularly in Europe among men having sex with men, whose extent is unprecedented. Since May 2022 we implemented MPX virus diagnosis by real-time PCR at university hospitals of Marseille, southern France. Here, we performed DNA metagenomic analyses of clinical samples from MPX virus-infected patients between June and July 2022, using next-generation sequencing with Illumina or Nanopore technologies. Twenty-five samples from 25 patients were studied. This allowed obtaining a MPX virus genome for 18 patients, essentially from genital skin lesions and rectal swabbing. All 18 genomes were classified in the MPX virus B.1 lineage, and we delineated five sublineages (A-E). We detected a high number of mutations (66-73) scattered along the MPX virus genomes relatively to the genome obtained from a human in Nigeria in 2018. Some non-synonymous mutations occurred in genes encoding central proteins, among which transcription factors and core and envelope proteins. They included two mutations that truncate RNA polymerase subunit RPO132 and a phospholipase D-like protein, which suggests gene inactivation. In addition, we identified that a large majority of nucleotide substitutions (94%) in the 18 MPX virus genomes were G>A or C>T, suggesting the action of human APOBEC3 enzymes. Finally, while we did not detect reads matching with main bacterial agents of sexually transmitted infections, >1,000 reads identified *Staphylococcus aureus* and *Streptococcus pyogenes* for 7 and 17 samples, respectively. These findings warrant a close genomic monitoring of MPX virus to get a better picture of this virus’ genetic evolution and mutational patterns, and they point out the common presence in monkeypox lesions of bacterial agents of skin superinfection, which warrants a close clinical monitoring in monkeypox patients.

## INTRODUCTION

Monkeypox virus (MPX virus) is an enveloped virus with a double-stranded DNA genome that is approximately 197,000 base pair (bp)-long and encodes for about 200 genes, half of which are reported to be involved in virion replication and morphogenesis while the other half comprises accessory genes involved in virus-host interactions (Senkevitch et al., 2021). This virus has an approximately 200 nm-large capsid that is complex and responsible for the brick shaped morphology of virions (Bahar et al., 2011). It is classified in family *Poxviridae*, subfamily *Chordopoxvirinae*, and genus *Orthopoxvirus*. According to the recently udpated viral taxonomy, it belongs to realm *Varidnaviria*, kingdom *Bamfordvirae*, phylum *Nucleocytoviricota*, class *Pokkesviricetes*, and order *Chitovirales* (https://www.ncbi.nlm.nih.gov/Taxonomy/Browser/wwwtax.cgi?id=10240; https://ictv.global/filebrowser/download/6719; https://ictv.global/taxonomy/taxondetails?taxnode_id=202104737). Due to the large size of the genome and virion, MPX virus had been earlier classified with the other poxviruses in a monophyletic group called nucleocytoplasmic large DNA virus (NCLDV) (Iyer et al., 2001), then in a proposed order named *Megavirales* that gathered NCLDV members and giant viruses isolated from amoebae (Colson et al., 2017).

MPX virus is endemic in humans in sub-Saharan Africa with two main foci of infections located in Central Africa and West Africa (Mauldin et al., 2022; Alakunle et al., 2022). Three viral clades named 1 to 3 have been delineated including clade 1 (or clade I according to the World Health Organization (WHO; https://www.who.int/news/item/12-08-2022-monkeypox--experts-give-virus-variants-new-names)) linked to outbreaks in Democratic Republic of Congo, clade 2 (or WHO clade IIa) linked to West Africa, and clade 3 (or WHO clade IIb) that encompasses viruses originating from the 2017-2018 outbreak that began in Nigeria and now those involved in the current 2022 multicountry outbreak (Berthet et al., 2021; Isidro et al., 2022b, Luna et al., 2022; Mauldin et al., 2022; Wang et al., 2022). Although human MPX virus infections imported from subSaharan Africa have occurred in non-endemic countries (Mauldin et al., 2022), they only caused sporadic cases or very limited outbreaks until 2022. An epidemic of MPX virus infections has arisen in May 2022 in non-endemic areas, the extent of which has greatly exceeded that of the previous epidemics (Vaughan et al., 2022; out et al., 2022). As of 12/09/2022, the number of confirmed cases was 57,315 worldwide and 3,785 in France (https://ourworldindata.org/monkeypox). The epidemiological and clinical features of this current epidemic indicate it is sexually-transmitted, which is new (Bragazzi et al., 2022).

The first MPX virus genome of this 2022 outbreak in non-endemic countries was released on 20/05/2022 in Portugal (Isidro et al., 2022a). In France, the first draft genome was released on 25/05/2022 in Southwest France (Croville et al., 2022). Phylogenomic studies of MPX viruses involved in this 2022 outbreak indicated that these viruses belong to clade 3 and likely have a single origin (Isidro et al., 2022b; Wang et al., 2022; Jones et al., 2022). They further indicated that these viruses comprise a lineage named B.1 that derives from the A.1 lineage whose members were associated with the large outbreak in 2017-2018 in Nigeria and with MPX virus exportation from Nigeria to the United Kingdom, Israel and Singapore in 2018-2019 (Mauldin et al., 2022; Isidro et al., 2022b). The B.1 lineage has been estimated to have emerged in Europe on early February 2022, most probably between November 2021 and May 2022 (Luna et al., 2022). Interestingly, the MPX viruses that cause the 2022 outbreak were found to diverge from those of the A.1 lineage by approximately 50 nucleotide substitutions (Isidro et al., 2022b; Wang et al., 2022; Jones et al., 2022), which is roughly 10 times more than expected on the basis of previous estimates of the orthopoxvirus mutation rate of 1–2 substitutions per genome per year (Firth et al., 2010).

Since May 2022 we implemented MPX virus diagnosis by real-time PCR (qPCR) at IHU Méditerranée Infection in Marseille, for the whole southeastern region (Provence-Alpes-Côte d’Azur) of France. Here we describe the 18 first MPX virus genomes and the bacterial sequences obtained from clinical samples of MPX virus-infected patients in the setting of clinical virology diagnosis.

## RESULTS

### Selection of Monkeypox virus-positive samples used for next-generation sequencing

Between the 22/05/2022 and the 22/07/2022 (two months), 307 patients were tested for the presence of MPX virus in the setting of clinical virology diagnosis. This was performed with an in-house real-time PCR assay (qPCR) (Scaramozzino et al., 2007) on several clinical samples including mostly skin lesions, and fecal and nasopharyngeal samples. On the basis of our experience of the yield of next-generation sequencing (NGS) of viral genomes (including SARS-CoV-2) (Colson et al., 2022) without targeted PCR amplification, NGS was performed for clinical samples with a cycle threshold value (Ct) of the diagnosis qPCR ≤25. Thus, 25 samples from 25 patients were used for NGS (Table 1). Their mean (± standard deviation) Ct was 22.5±1.7 (range, 20.0-25.0). They were skin (n= 14; 56%), rectal (n= 10; 40%), and pharyngeal (n= 1; 4%) samples collected by swabbing. They were collected between the 13/06/2022 and the 22/07/2022.

**Table 1.** Main features of patients and samples.

| Sampling date | Sample type | qPCR<br>orthopoxvirus (Ct) | Number of reads | Number of reads<br>mapped on the<br>genome GenBank<br>Accession no.<br>ON563414.3 | Mean<br>sequencing<br>depth | Genome<br>coverage | Monkeypox virus clade<br>3 lineage | Genome identifier |
| --- | --- | --- | --- | --- | --- | --- | --- | --- |
| 13/06/2022 | Skin lesion | 21 | 1 181 088 | 2 461 | 93 | 100,00 | B.1 | hMPXV-IHU00001 |
| 01/07/2022 | Rectal sample | 22 | 1 694 878 | 148 297 | 166 | 99,98 | B.1.1 | hMPXV-IHU00002 |
| 04/07/2022 | Skin lesion | 20 | 1 880 874 | 57 666 | 60 | 99,98 | B.1 | hMPXV-IHU00003 |
| 04/07/2022 | Skin lesion | 25 | 1 795 572 | 50 310 | 58 | 99,98 | B.1 | hMPXV-IHU00004 |
| 04/07/2022 | Rectal sample | 21 | 1 470 090 | 60 224 | 65 | 99,98 | B.1 | hMPXV-IHU00005 |
| 04/07/2022 | Rectal sample | 23 | 1 073 538 | 11 197 | 12 | 96,55 | B.1 | hMPXV-IHU00006 |
| 05/07/2022 | Skin lesion | 22 | 2 228 394 | 19 577 | 17 | 93,43 | B.1 | hMPXV-IHU00008 |
| 06/07/2022 | Rectal sample | 22 | 1 531 952 | 66 461 | 67 | 99,96 | B.1 | hMPXV-IHU00007 |
| 07/07/2022 | Rectal sample | 24 | 3 125 930 | 9 762 | 8 | 73,50 | B.1 | - |
| 07/07/2022 | Rectal sample | 24 | 1 221 748 | 3 989 | 3 | 34,31 | / | - |
| 07/07/2022 | Skin lesion | 24 | 2 000 646 | 85 249 | 101 | 99,98 | B.1 | hMPXV-IHU00009 |
| 18/07/2022 | Skin lesion | 23 | 1 654 956 | 1 195 | 1 | 4,21 | / | - |
| 18/07/2022 | Rectal sample | 22 | 1 030 066 | 3 311 | 3 | 31,74 | / | - |
| 18/07/2022 | Rectal sample | 20 | 1 577 758 | 38 162 | 32 | 98,42 | B.1 | hMPXV-IHU00010 |
| 19/07/2022 | Skin lesion | 25 | 1 301 248 | 31 687 | 32 | 99,37 | B.1.1 | hMPXV-IHU00011 |
| 19/07/2022 | Skin lesion | 20 | 3 234 274 | 59 725 | 46 | 99,59 | B.1 | hMPXV-IHU00012 |
| 20/07/2022 | Skin lesion | 25 | 4 957 512 | 16 273 | 11 | 87,12 | B.1 | - |
| 20/07/2022 | Rectal sample | 25 | 1 648 064 | 3 671 | 3 | 34,69 | / | - |
| 20/07/2022 | Skin lesion | 22 | 5 592 602 | 59 992 | 38 | 99,53 | B.1 | hMPXV-IHU00013 |
| 21/07/2022 | Skin lesion | 24 | 886 636 | 2 186 | 1 | 13,69 | / | - |
| 22/07/2022 | Skin lesion | 21 | 4 506 848 | 35 232 | 25 | 99,13 | B.1 | hMPXV-IHU00014 |
| 22/07/2022 | Skin lesion | 23 | 1 967 452 | 18 610 | 12 | 90,41 | B.1 | hMPXV-IHU00015 |
| 22/07/2022 | Nasopharyngeal sample | 23 | 5 977 822 | 19 802 | 13 | 96,69 | B.1 | hMPXV-IHU00016 |
| 22/07/2022 | Skin lesion | 21 | 4 882 696 | 22 155 | 15 | 97,90 | B.1 | hMPXV-IHU00017 |
| 22/07/2022 | Rectal sample | 21 | 3 440 078 | 16 259 | 11 | 93,80 | B.1 | hMPXV-IHU00018 |

### Yield of next-generation sequencing and obtaining of full-length Monkeypox virus genomes

All but one NGS were performed by the Illumina technology on a MiSeq instrument (Illumina Inc., San Diego, CA, USA); a single sample was used for NGS with the Nanopore technology on a GridION instrument (Oxford Nanopore Technologies Ltd., Oxford, United Kingdom). The mean number of NGS reads per sample was 2,474,509 (range, 886,636-5,977,822) (Table 1). The mean number of reads per sample identified as from MPX virus genomes, due to their mapping on the genome GenBank Accession no. ON563414.3 that was obtained from a human specimen collected in May 2022 in the USA and is used as a B.1 lineage reference by Nextclade (https://nextstrain.org/monkeypox/hmpxv1) (Aksamentov et al., 2021), was 33,738 (range, 1,195-148,297) (Table 1). Mean genome coverage relatively to the genome GenBank Accession no. ON563414.3 was 81.8±30.7% (range, 4.2-100.0%). Mean sequencing depth for mapping was 36 reads (range, 1-166 reads). These results were similar when excluding from the analysis the genome generated by the Nanopore technology, for which 1,181,088 NGS reads have been generated, 2,461 reads were identified as from a MPX virus genome, sequencing depth was 93.4 and genome coverage was 100.0% (Table 1).

A total of 18 near full-length (coverage of at least 90%) or full-length genomes were obtained and further characterized. For these 18 genomes, the mean number of reads mapped on the genome GenBank Accession no. ON563414.3 per sample was 44,615 (range, 1,073,538-5,977,822) (Table 1). Mean sequencing depth was 48±41 reads (range, 11-166 reads), and mean genome coverage was 98.0±2.8% (range, 90.4-100.0%). These 18 genomes were obtained from 18 different patients. Eleven were obtained from skin lesions including 5 localized on the penis, 6 were obtained from rectal samples, and one was obtained from a pharyngeal sample. These samples were collected between the 13^rd^ of June and the 22^st^ of July 2022. The mean Ct of the qPCR used for MPX virus diagnosis was 22.0±1.6 (20.0-25.0).

### Characteristics of the Monkeypox virus genomes

All 18 MPX virus genomes obtained in the present study (GenBank Accession no. OP382478-OP382495) are classified in the B.1 lineage of clade 3 (or clade IIb according to the WHO classification), and two are classified into the B.1.1 sublineage (https://nextstrain.org/monkeypox/hmpxv1) (Aksamentov et al., 2021) (Table 1). Relatively to the MPX virus genome ON563414.3 of the B.1 lineage, 6 of the 18 genomes are genetically identical while 12 harbor at least one mutation (Tables 2, 3, 4). These 12 latter genomes harbor a mean number of 3.3±2.2 mutations (range, 1-7 mutations) relatively to ON563414.3. Mean nucleotide identity between these genomes calculated by pairwise comparison is 99.8±0.2% (range, 99.3-100.0%).

**Table 2.**
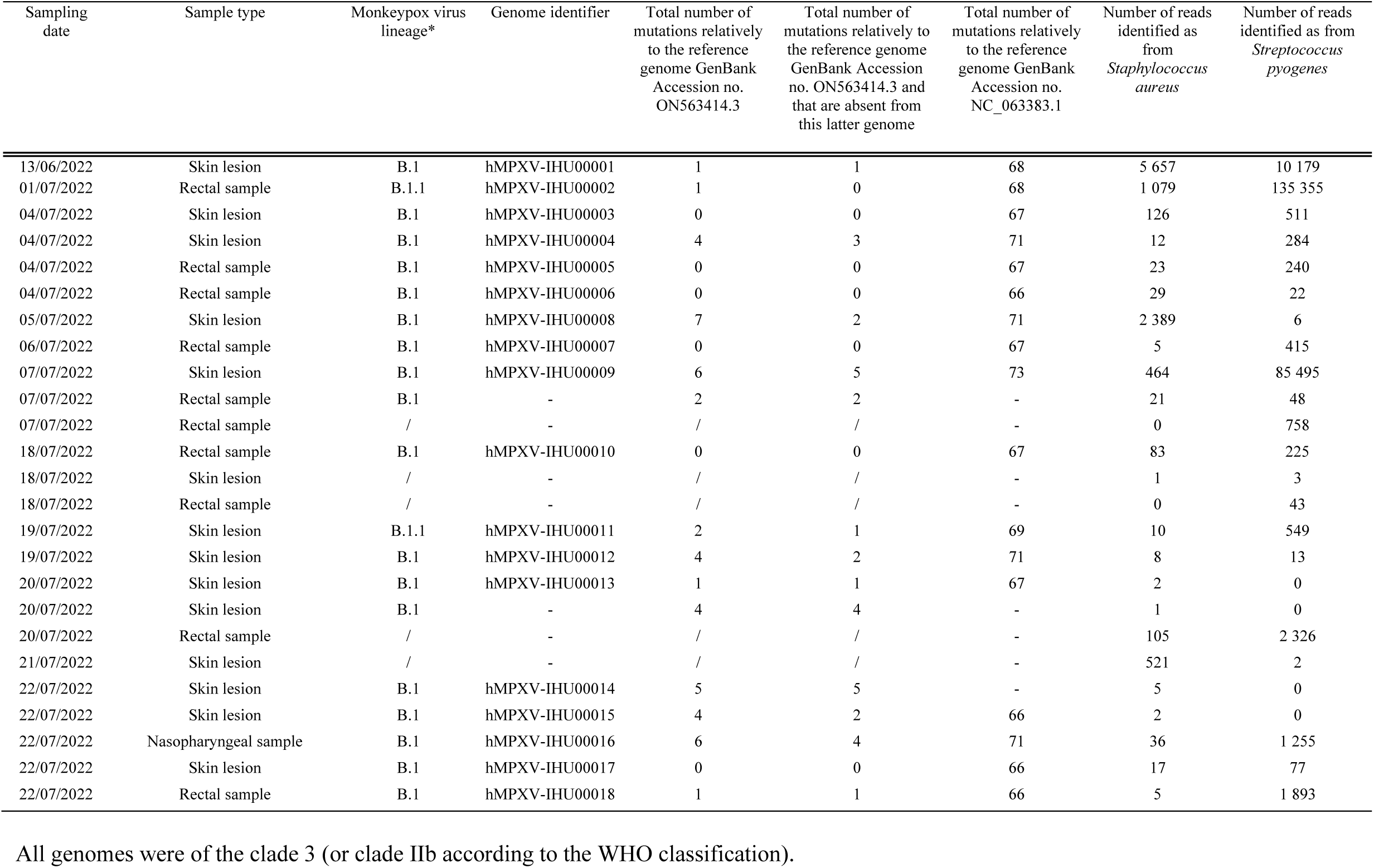
Main features of Monkeypox virus genomes.

**Table 3.**
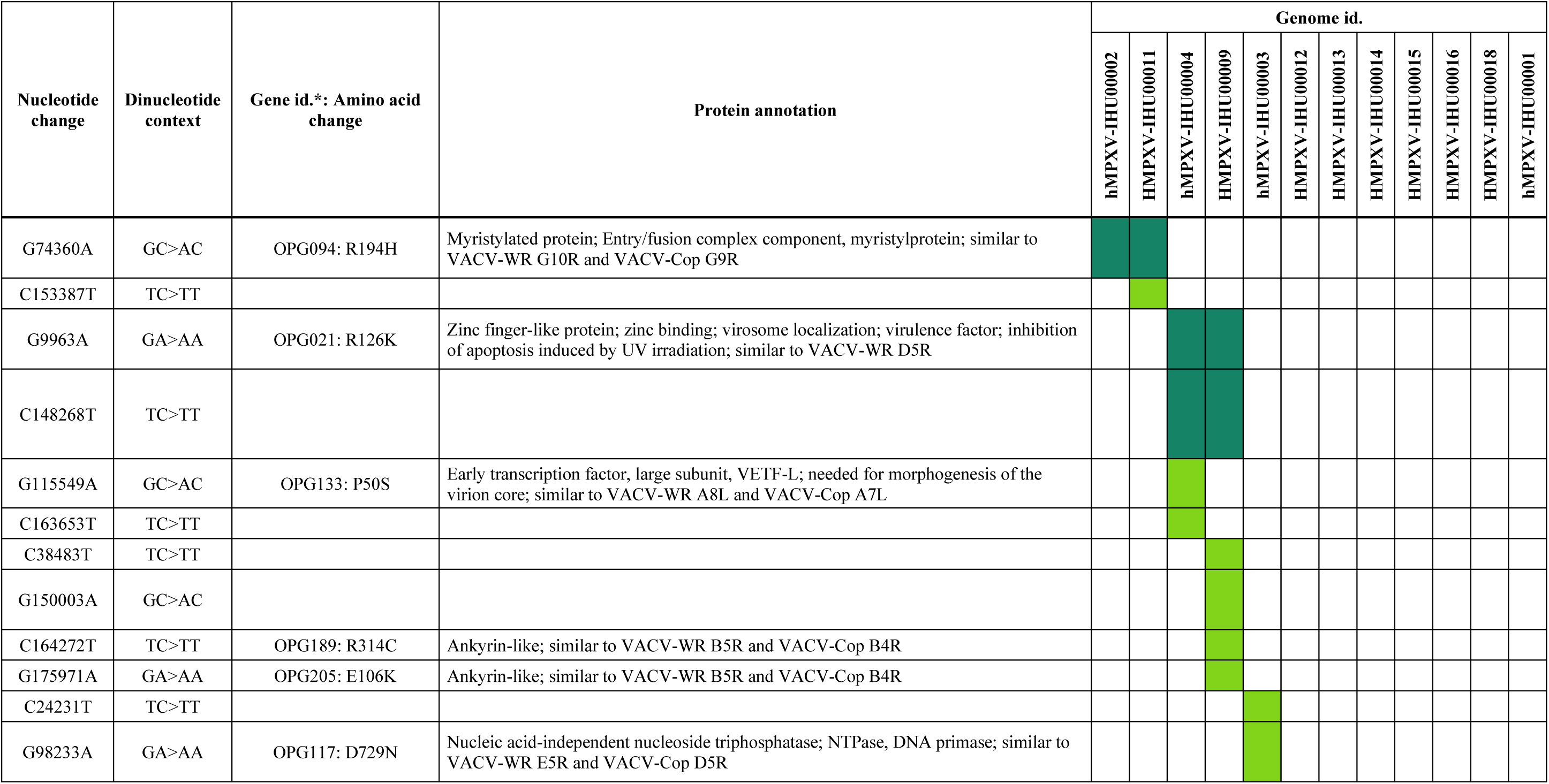
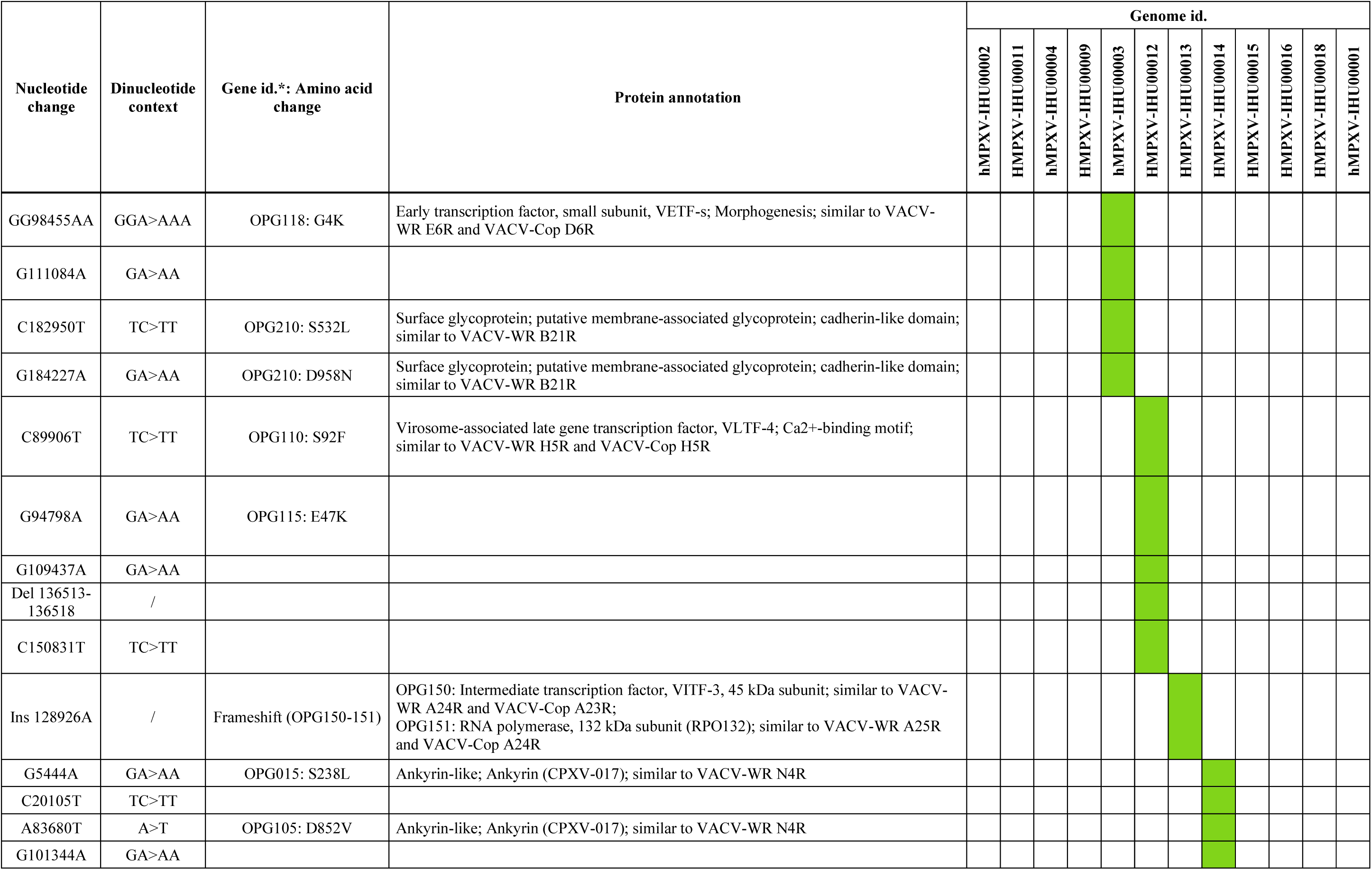
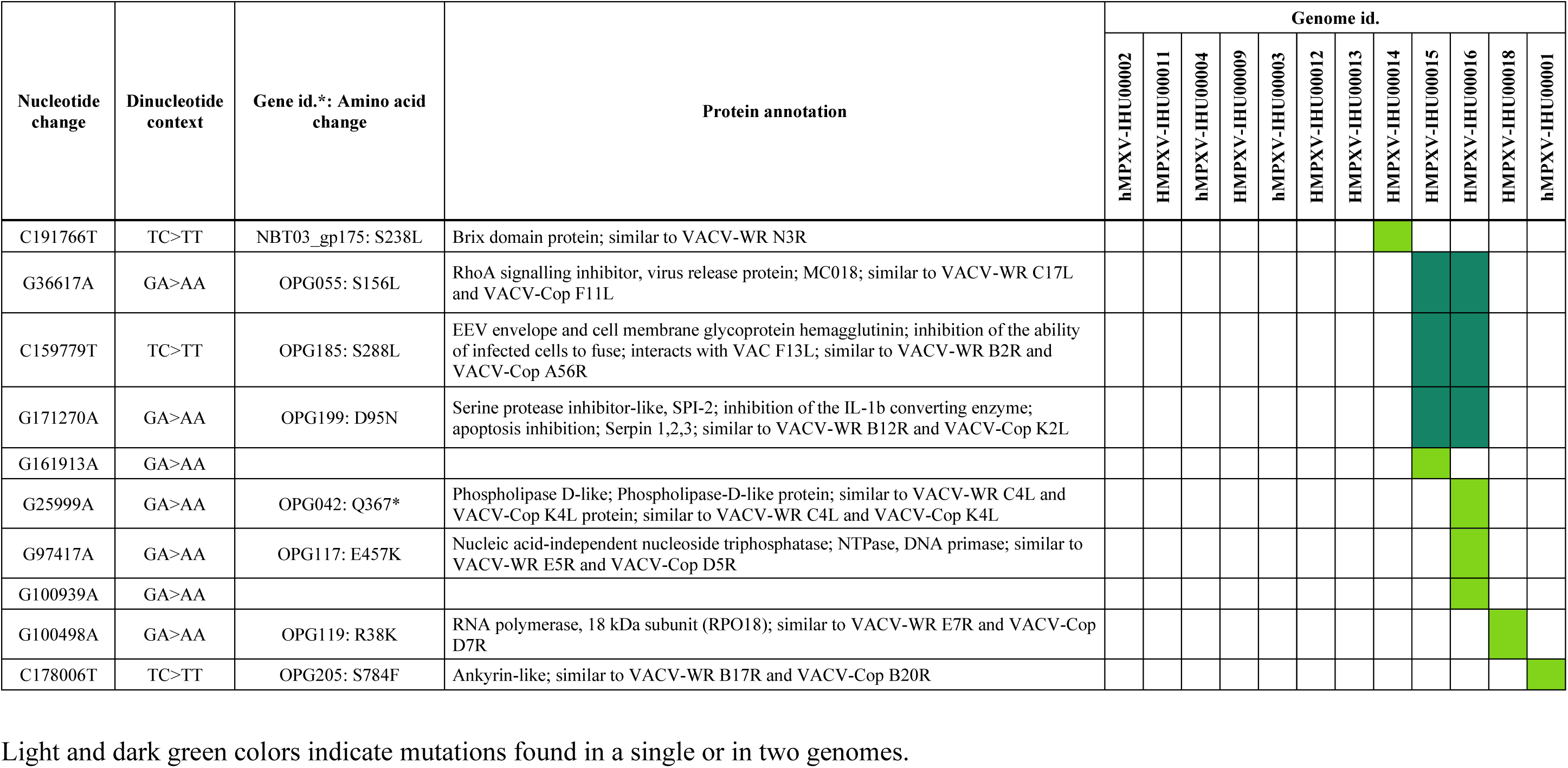
Nucleotide and amino acid change in Monkeypox virus genomes obtained in the present study relatively to genome GenBank Accession no. ON563414.3, for the genomes that harbor at least one mutation.

**Table 4.**
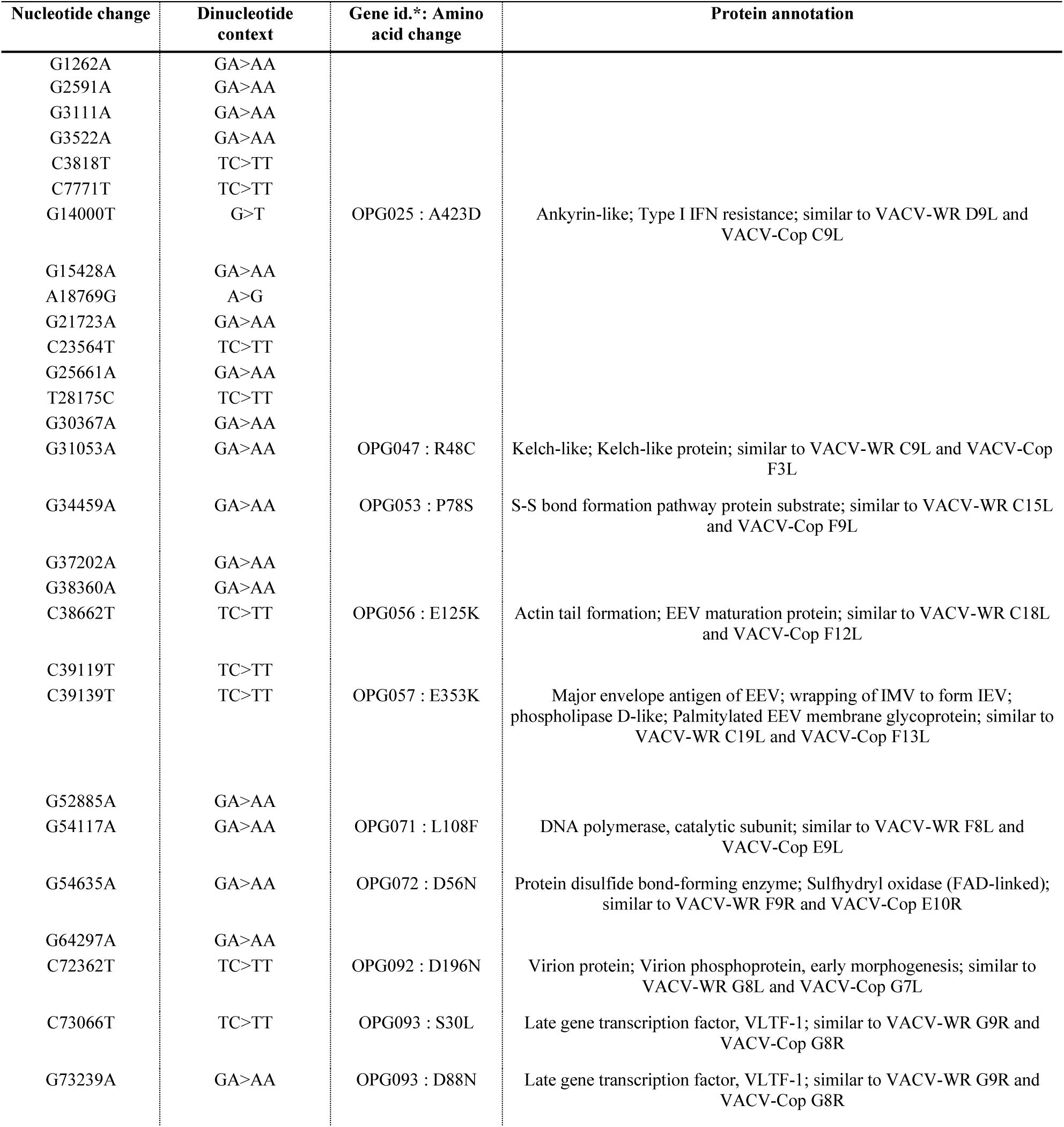
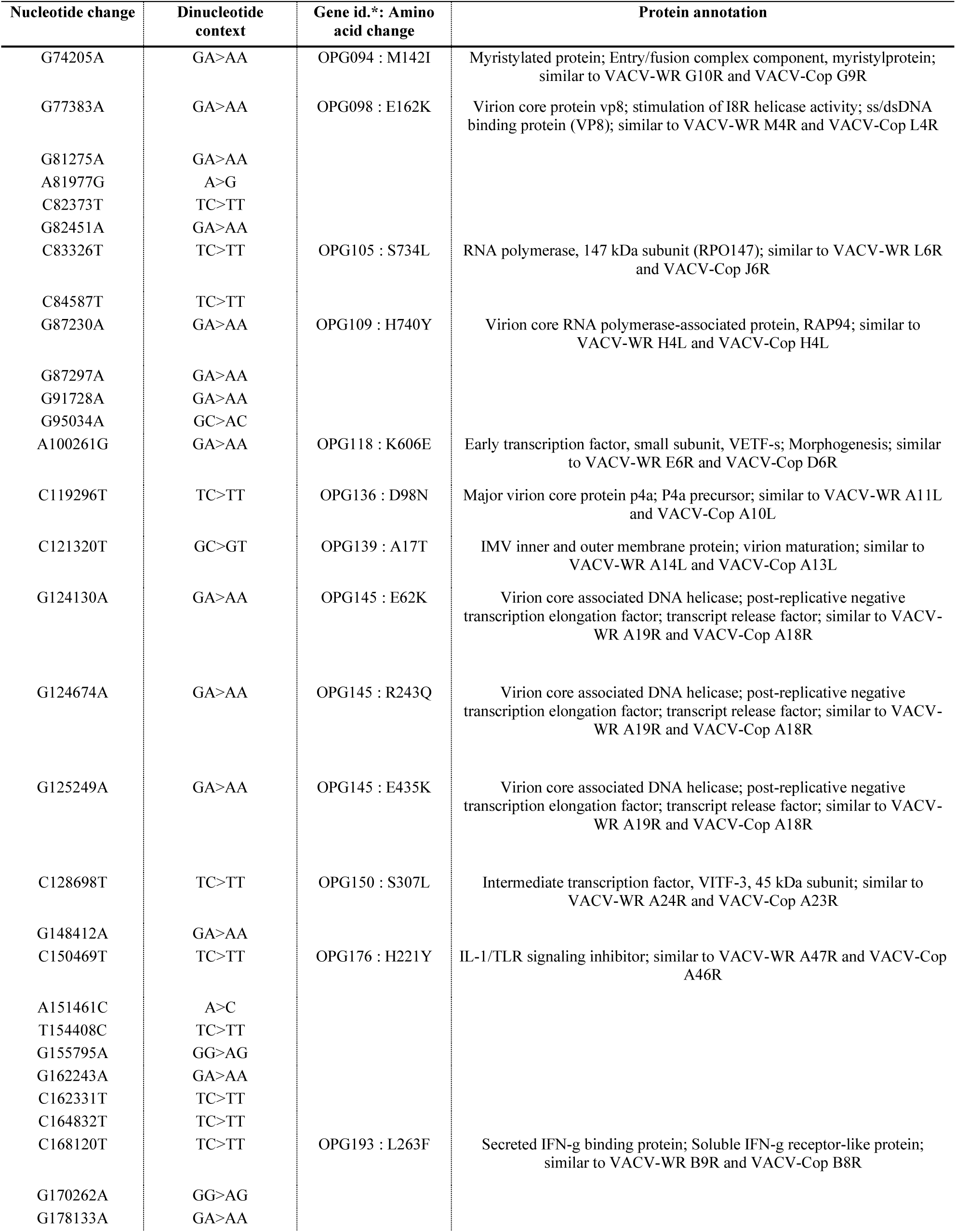
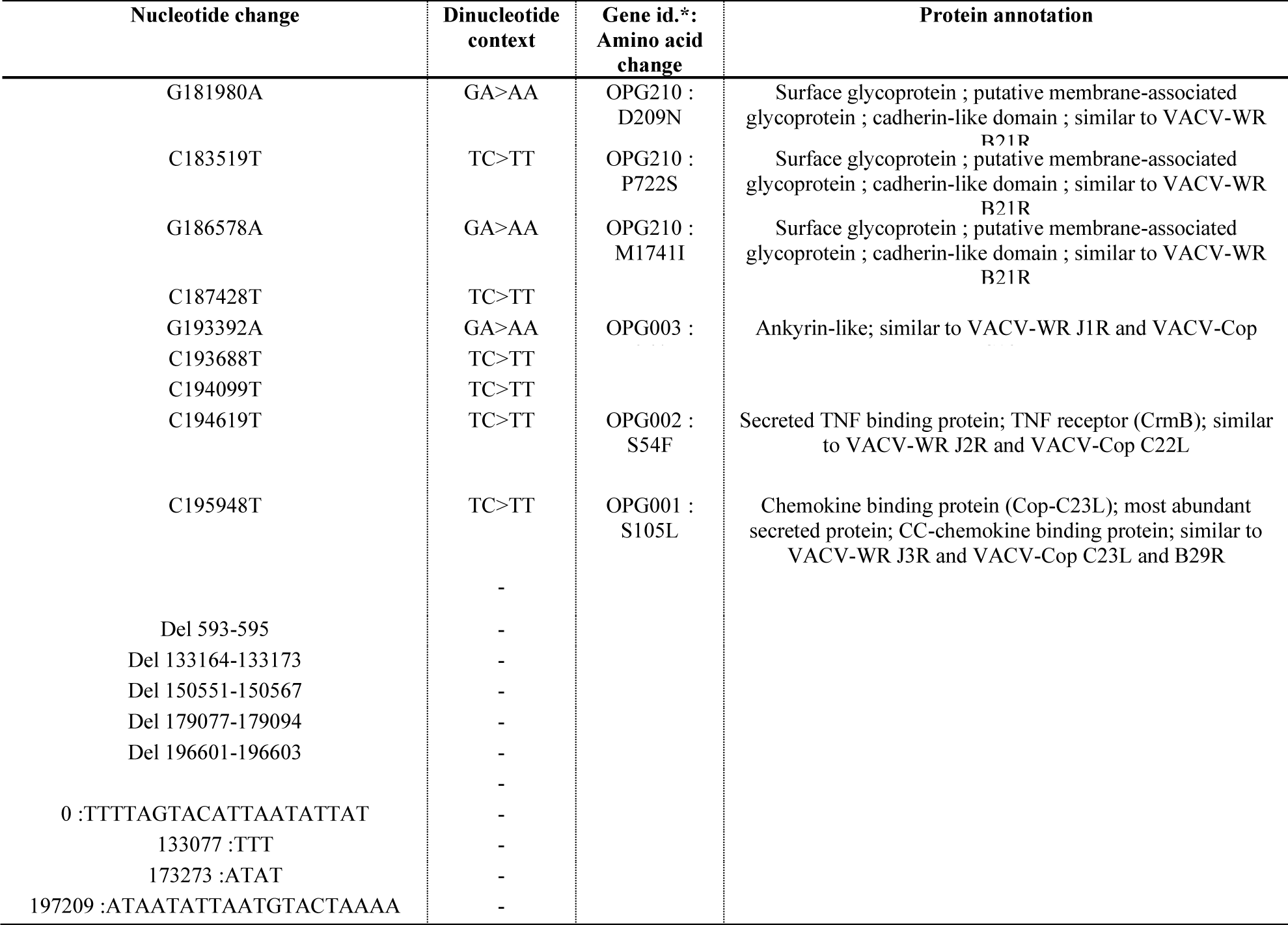
Nucleotide and amino acid change in Monkeypox virus genomes obtained in the present study and of the B.1 lineage relatively to genome GenBank Accession no. NC_063383.1. Mutations specific of the genomes obtained in the present study relatively to genome GenBank Accession no. NC_063383.1 are shown in Figure 1.

A total of 36 different mutations are present in at least one of the 18 viral genomes, and each of them are present in one or two genomes. These mutations are non-synonymous in 22 cases and synonymous in 14 cases. The non-synonymous mutations are located in 20 genes, being scattered along the viral genome (Tables 3, 4; Supplementary Figure S1). These 20 genes notably include two RNA polymerase subunits; two early transcription factor subunits and one intermediate and one late transcription factors; a nucleoside triphosphatase; and four ankyrin-repeat containing proteins. Strikingly, one insertion generates in one MPX virus genome (hMPXV-IHU00013; GenBank Accession no. OP382490.1) a frameshift that is predicted to lead to a truncation of the RNA polymerase subunit RPO132. For this genome, 47 NGS reads harbor the insertion while a single one does not (Supplementary Figure S2). In addition, this insertion was found for only one of 17 genomes sequenced with the Illumina technology although NGS of the 16 other genomes was performed into three different NGS runs on the Illumina MiSeq instrument. Moreover, the insertion is also present in another B.1 lineage genome obtained from a patient sampled in Portugal in July 2022 (GISAID Accession no. EPI_ISL_14515228). Therefore, these data suggest it is unlikely that this mutation was a technical artefact. Otherwise, one substitution in the genome hMPXV-IHU00016 (GenBank Accession no. OP382493.1) was detected that generates a stop codon and is predicted to truncate a phospholipase D-like protein.

Relatively to the genome GenBank Accession no. NC_063383.1 (isolate: MPXV-M5312_HM12_Rivers-001) that that belongs to MPKV clade 2 (or clade IIa according to the WHO classification) and was obtained from a human specimen collected in Nigeria in August 2018 (Mauldin et al., 2022), the 18 genomes obtained in the present study harbor a mean number of 68.3±2.3 mutations (range, 66-73 mutations) (Table 2). The total number of nucleotide substitutions is 67. These include 29 non-synonymous mutations leading to amino acid changes in 24 proteins. Some of these proteins are predicted to be involved in virion replication or to be important virion structural components, such as DNA polymerase and RNA polymerase subunits (OPG071, OPG105; in reference to the gene repertoire of genome NC_063383.1); early and late transcription factors (OPG118, OPG93); virion core proteins (OPG098, OPG136); or the major envelope protein (OPG057). Also, three amino acid substitutions are detected in proteins annotated as a putative membrane-associated glycoprotein (OPG210) and a virion core associated DNA helicase suspected to be a post-replicative negative transcription elongation factor (OPG145).

We identified that a large majority of nucleotide substitutions (94%) are G>A (56%) and C>T (36%), which can be signatures of the action of human apolipoprotein B mRNA-editing catalytic polypeptide-like 3 (APOBEC3) enzymes. As the proportion of A, C, G and T nucleotides in the MPX virus genomes are approximately 34%, 17%, 16% and 33%, respectively, which corresponds to a GC-content of 33%, there is therefore a strong mutation bias. In addition, 51 of 57 G>A substitutions are in a GA dinucleotide context (i.e., changes are GA>AA; 4 G>A substitutions are in a GC context and 2 in a GG dinucleotide context), while 38 of 39 C>T substitutions are in a TC context (i.e., changes are TC>TT), which are further hints that these nucleotide changes might be caused by the deaminase activity of APOBEC3 enzymes (McDaniel et al., 2020; Isidro et al., 2022b) (Tables 3, 4).

### Clusters of Monkeypox virus genomes based on mutation patterns and phylogenetic analyses

We searched for the MPX virus genomes the most similar to those obtained in the present study based on mutation patterns and phylogenetic analyses (Figure 1; Supplementary Table S1). We could delineate five sublineages (we named A to E) that involve either one (in two cases) or two (in three cases) genomes from our laboratory. Sublineage D corresponds to the described sublineage B.1.1 (https://nextstrain.org/monkeypox/hmpxv1) (Aksamentov et al., 2021). No pair of genomes identical between each other was identified. These five lineages encompass genomes sharing all or some of the mutations identified relatively to genome ON563414.3 in the present work (Supplementary Table S1). The two genomes of sublineage D share one mutation, the two genomes of sublineage E share two mutations and the two genomes of sublineage A share three mutations. In addition, these sublineages are supported by bootstrap values comprised between 81 and 100% in the phylogeny reconstruction (Figure 1). Ten genomes obtained here were not clustered neither together nor with their most similar counterpart genomes. The MPX virus genomes the most closely related to those obtained in our laboratory were obtained in various countries including Germany (sublineages A and D); Canada, USA, and Brazil (sublineage B); United Kingdom and Finland (sublineage C); United Kingdom, Germany and France (sublineage E).

**Figure 1.**
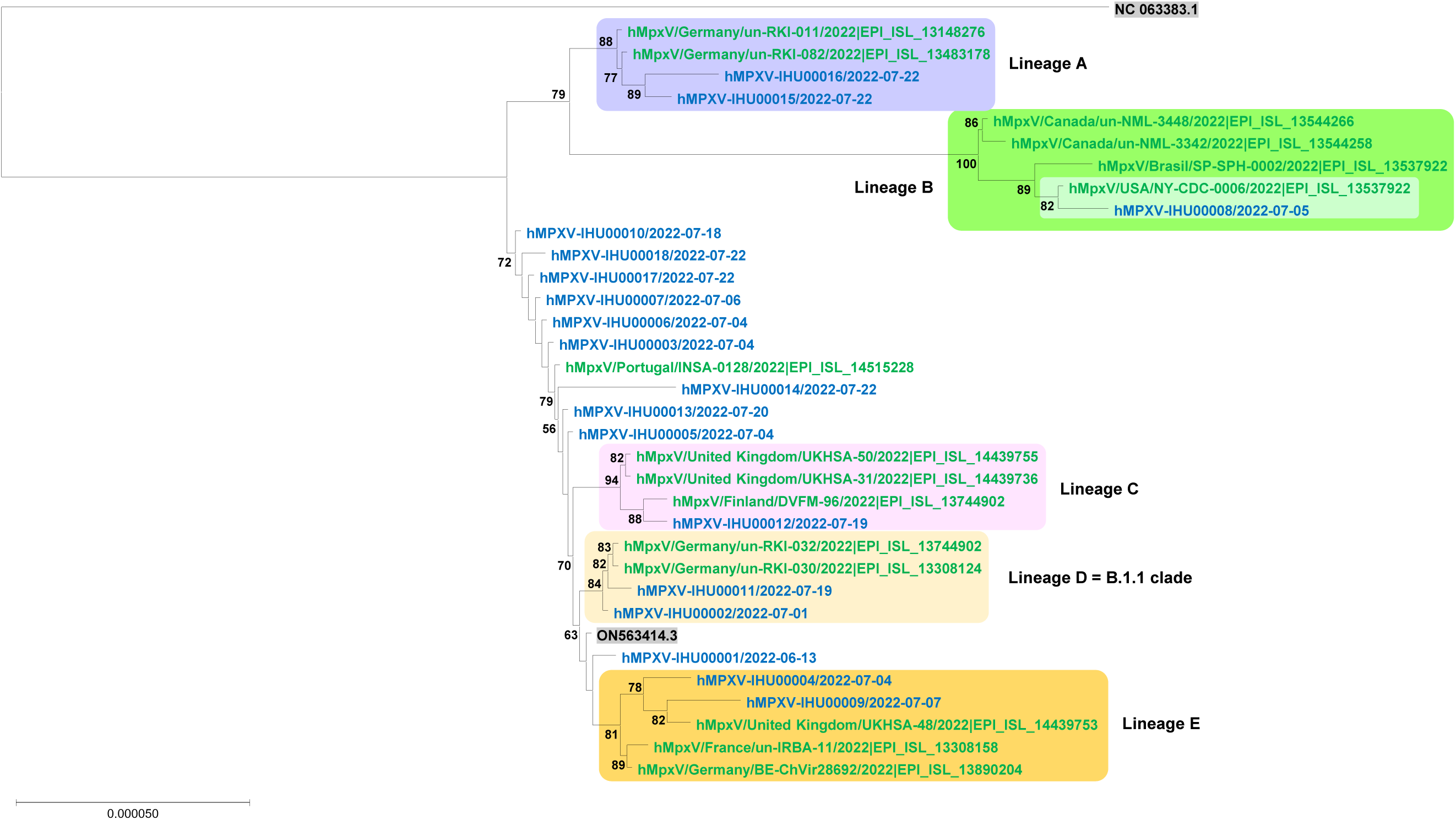
Phylogenetic analysis based on Monkeypox virus genomes obtained in the present study and genomes that are those the most similar to them. For Information on the genomes from the GISAD sequence database (https://gisaid.org/), see Supplementary Table S3.

### Monkeypox virus genome annotation

We attempted to annotate the first MPX virus genome (hMPXV-IHU00001; GenBank Accession no. OP382478.1) that we obtained using GeneMarkS (Besemer et al., 2001). Of the 208 gene products predicted in hMPXV-IHU00001, 124 are strictly identical in length and amino acid composition with those of NC_063383 (Supplementary Table S2). Of the 61 remaining gene products, 33 showed at least 99% identity in length and amino acid composition. In 6 cases, these gene products had paralogs. A total of 22 predicted proteins had no significant matches in the set of the NC_063383 predicted proteins. Of them, 20 had significant matches with predicted proteins of other poxviruses, of MPX virus in 17 cases, and of Cowpox virus or Vaccinia virus in 4 cases. Five of these predicted proteins were equal in size or larger than 100 amino acids, and they are annotated as a virosome component, two cowpox A-type inclusion protein, an interleukine-1-binding protein, or a TNF-alpha receptor-like protein. Conversely, for all 179 predicted proteins of NC_063383, an ortholog was found in hMPXV-IHU00001.

### Metagenomic analysis of next-generation sequencing runs

As NGS on the 25 clinical samples from the 25 patients was performed in the absence of prior MPX virus specific PCR amplification, we analyzed sequencing runs as a DNA metagenomes using Kraken2 (Wood et al., 2014). We did not detect NGS reads matching with bacteria responsible of sexually transmitted infection including *Chlamydia trachomatis*, *Neisseria gonorrhoeae*, or *Treponema pallidum*. In contrast, we detected reads identifying *Staphylococcus aureus* and *Streptococcus pyogenes*, two agents of skin lesion superinfection (Table 2; Supplementary Figure S3). The mean number of reads was 424 (range, 0-5,657) and 9,588 (0-135,355) for these bacteria, respectively. There were between 10 and 100 NGS reads identifying *S. aureus* for 7 samples, between 100 and 1,000 reads for 4 samples, and >1,000 reads for 3 samples. The 7 samples from which >100 reads were generated were 5 skin lesions including 4 localized on the penis, and 2 rectal samples. Regarding reads identifying *S. pyogenes*, there were between 10 and 100 NGS reads for 5 samples, between 100-1,000 reads for 7 samples, and >1,000 reads for 6 samples. The 13 samples from which >100 reads were generated were 5 skin lesions, 7 rectal samples, and one nasopharyngeal samples.

## DISCUSSION

Here, we obtained 18 MPX virus genomes, which increases by four times the number of genomes available for France as of 09/09/2022. At the global level, 2,117 genomes were released in GISAID (https://gisaid.org/) (Elbe et al., 2017) at that time. All these genomes were found to belong to the B.1 lineage, as for the case of other viral genomes released for the 2022 outbreak (Wang et al., 2022; Isidro et al., 2022b). In addition, we identified that two genomes belong to the B.1.1 sublineage, and identified 4 other lineages encompassing one or two viral genomes obtained in our laboratory. The phylogenomic analyses of the 18 genomes indicate firstly that MPX virus detected among patients tested in our institute did not comprise a particular lineage, apart from other viruses involved in the 2022 outbreak in non-endemic areas. Second, they show that the genomes the most closely related to those obtained in our laboratory are from Europe and North and South America. A single other genome among those most closely related to ours was from France (un-IRBA-11/2022|EPI_ISL_13308158); however, this must be interpreted with caution by considering that only 6 genomes have been released to date by French laboratories according to the GISAID sequence database as of 09/09/2022.

The analysis of the 18 MPX virus genomes obtained in the present study showed the presence of non-synonymous mutations leading to amino acid changes in dozens of viral proteins involved in virion replication and morphogenesis, as well as in virus-host interactions, including some that are deemed to be central for the viral cycle. Such mutations have been detected in several other European studies (Isidro et al., 2022b; Wang et al., 2022; Jones et al., 2022; Otoole and Rambaut, 2022). Proteins affected by amino acid changes include for instance DNA and RNA polymerase subunits; transcription or elongation factors; virion core proteins; the major envelope protein; a putative membrane-associated glycoprotein; ankyrin-repeat containing proteins; and a phospholipase D-like protein. The impact of these mutations on the function of these proteins and on the viral replication cycle is worthy to be investigated. Interestingly, we observed, as previously reported by other teams (Isidro et al., 2022b; Wang et al., 2022; Jones et al., 2022; Otoole and Rambaut, 2022), that three amino acid changes implicated the B21 protein (OPG210 gene in the genome NC_063383.1), a surface putative membrane-associated glycoprotein similar to Vaccinia virus VACV-WR B21R protein. This protein has been found to be highly immunogenic (Hammarlund et al., 2005), and this questions if these amino acid changes may have been selected as they might lead to viral immune escape. Regarding ankyrin-repeat containing proteins, they are known to be involved in a broad range of protein-protein interactions including in virus-host interaction (Senkevitch et al., 2021; Al-Khodor et al., 2010; Pagnier et al., 2015). The genomes of NCLDV, including poxviruses, have been described to encode multiple ankyrin repeats. Ankyrin-repeat containing proteins were proposed in poxviruses to be capable of interacting with the ubiquitin-proteasome system to modulate cellular processes (Barry et al., 2010). The cases of nucleotide changes in genes encoding the phospholipase-D and a RNA polymerase subunit are particularly worthy of interest. Indeed, they may lead to truncated proteins. The phospholipase-D (encoded in MPX virus genomes by the ortholog of genes named OPG042 in MPX virus genome NC_063383.1 or K4L in Vaccinia virus) is predicted to be a K4 endonuclease, and to be nonessential as the deletion of its gene was reported to not affect Vaccinia virus replication or virulence (Selvy et al., 2011). The presence of a frameshift in the gene encoding RNA polymerase subunit RPO132 is particularly intriguing. Indeed, unlike many other DNA viruses, poxviruses replicate in the host cell cytoplasm within viral factories, and their genomes encodes for their own replication machinery. Of note, for a giant DNA virus of the family *Marseilleviridae*, it was reported that the initial stage of replication cycle included the recruitment to the cytoplasmic viral factory of the transcription machinery of the amoebal host nucleus (Fabre et al., 2017); this suggests that, in some cases, the viral RNA polymerase might be dispensable at some stages of viral replication. Beyond, gene loss is a known trait of poxvirus evolution, and it was notably suspected to be associated with a restricted host range but also with an efficient human-to-human spread in orthopoxviruses (Senkevitch et al., 2021; Upton et al., 2003; Hendrickson et al., 2010). In addition, it was observed in MPX virus genomes recovered from 10 of 60 specimens collected from humans between 2005 and 2007 in the Democratic Republic of the Congo (Kugelman et al., 2014). This was the consequence of a deletion in the gene encoding an orthopoxvirus major histocompatibility complex class I-like protein, and was reported as to seemingly correlate with human-to-human transmission.

Another interesting observation is the high number of mutations (between 66 and 73) in MPX virus genomes obtained here compared to the NC_063383.1 genome obtained from a human specimen collected in Nigeria in August 2018, which is in the same range than previously reported (Isidro et al., 2022b; Wang et al., 2022; Jones et al., 2022; Otoole and Rambaut, 2022). This corresponds to a greater mutation rate than that expected based on previous assessment for orthopoxviruses that was estimated to be 1–2 nucleotide substitutions per genome per year (i.e. in the approximate range of 10^-6^-10^-5^ substitutions per site per year) (Firth et al., 2010; Babkin and Babkina, 2011), and might represent accelerated evolution. This could be related to the action of APOBEC3 enzymes, which are cytidine deaminases that acts on single-stranded DNA during the replication or transcription of double-stranded DNA (Harris et al., 2004; Armitage et al., 2008). As previously reported, we identified a strong mutation bias with a large majority of mutations (94%) being G>A (57%) and C>T (37%), suggesting that they could have occurred because of genome editing by human APOBEC3 enzymes, which perform nucleotide deamination. Indeed, these cellular enzymes may have promoted an excess of mutations in the MPX virus genomes and might have driven their short-term evolution (O’Toole and Rambaut, 2022; Isidro et al., 2022b; Jones et al., 2022). Interestingly, a human APOBEC3 enzyme, APOBEC3A, was reported to be expressed in keratinocytes and to generate G>A and C>T substitutions in human papillomavirus sequences (Vartanian et al., 2008). The high mutation rate in the MPX virus genomes may also be the consequence of significant spread by sexual transmission in the MSM community, as this was the case in 2017 for the hepatitis A virus in Europe, with an outbreak triggered by a pride gathering (Freidl et al., 2017). Whatever the causes of their generation, these numerous mutations may have led to modifications in some viral proteins impacting the tropism, the transmissibility, or the pathogenicity of MPX virus.

Finally, another interesting finding of the present study is the presence of NGS reads identifying sequences from *S. aureus* and *S. pyogenes* in a majority of MPX virus-positive clinical specimens, particularly skin lesions. To our knowledge, such data are reported for the first time on the basis of metagenomic analyses. Secondary bacterial infections have been reported as occurring in the setting of monkeypox in 6% of 34 cases in Italy (Moschese et al., 2022), and in 48% of 40 cases in Nigeria and up to 100% of the HIV-positive individuals (Ogoina et al., 2020) They were reported to be among the main causes of hospitalization with severe perianal pain (Moschese et al., 2022). Cellulitis was reported in 11% of cases in UK (Girometti et al., 2022). In the study conducted in Italy, one patient presented a penis cellulitis documented by *S. aureus* and *S. pyogenes* culture, which required prolonged antibiotic therapy (Moschese et al., 2022). These findings suggest that these two bacteria may cause superinfections but further studies are needed to elucidate the real prevalence of bacteria in association with MPX virus lesions. Although secondary bacterial infection of skin lesions are recognized as a common complication of monkeypox, the WHO recommends not to use antibiotics therapy or prophylaxis in patients with uncomplicated monkeypox because of the risk of emergence of multidrug-resistant bacteria and of potential side-effects such as *Clostridioides difficile* diarrhoea (WHO, 2022). WHO only recommends a monitoring of skin lesions for superinfection associated with cellulitis or abscess and in case of superinfection to treat by antibiotics that are active against methicillin-sensitive *S. aureus* and *S. pyogenes*. Regarding US Centers for Disease Control and Prevention (CDC) guidelines, they recommend that antibiotic treatment should be considered in people who have secondary bacterial skin infections (CDC, 2022). In addition, on a total of 14 guidelines on monkeypox clinical management released worldwide until 2021, only two were considering, and recommending antibiotherapy for secondary complications (Webb et al., 2022). The first one was published by the Chinese Ministry of Health and the second one by the Nigeria Centre for Disease Control. Recently, Fournier’s gangrene was reported in London in a 47-year-old man whose penile swab sampling allowed culturing *S. aureus* and *Streptococcus dysgalactiae* (Patel et al., 2022). In the current MPX outbreak, the morbidity linked to bacterial superinfection of skin lesions warrant their close monitoring, to allow a prompt administration of antibiotics in case of bacterial superinfection.

Overall, previous findings warrant a close genomic monitoring of MPX virus to get a better picture of the genetic evolution and mutational patterns of this virus that came into the light in non-endemic countries with the 2022 outbreak and have been revealed to spread globally (Alakunle et al., 2022). The role of gene losses in MPX virus transmissibility and replication, and that of APOBEC3 enzymes in the increased mutation rate observed in MPX virus genomes also deserves particular attention as their expression and deaminase activity can be modulated by various infections and has been linked to various cancers (Harris et al., 2004; Vartanian et al., 2008). Finally, the present study points out the detection of bacterial agents of skin superinfections concomitantly with the sequencing and characterization of MPX virus genomes, which warrants a close monitoring of such potential superinfections in monkeypox patients.

## MATERIALS AND METHODS

### Diagnosis of Monkeypox virus infections

Orthopoxvirus DNA was detected by a qPCR assay as previously described (Scaramozzino et al., 2007), with a T4 phage DNA internal control (Ninove et al., 2011), using the LightCycler Multiplex DNA Virus Master kit (Roche Diagnostics, Mannheim, Germany) on a LightCycler 480 instrument (Roche Diagnostics).

### Next-generation Monkeypox virus genome sequencing

NGS of MPX virus genomes was performed after we could obtain the authorization of the National Agency for the Safety of Medicines and Health Products (ANSM) (detention authorization number ADE-145582022-3 and manipulation authorization number AMO-145592022-5) to work on MPX virus nucleic acids. Based on our experience of the yield of NGS of viral genomes (including SARS-CoV-2) (Colson et al., 2022) in absence of prior targeted PCR amplification, we chose to perform NGS for clinical samples with a cycle threshold value (Ct) of the diagnosis qPCR ≤25. Selected samples from MPX virus-positive patients were directly sequenced without prior PCR amplification by the Illumina technology with the Nextera XT paired-end strategy on a MiSeq instrument (Illumina Inc., San Diego, CA, USA) or, in one case, with the Nanopore technology on a GridION instrument (Oxford Nanopore Technologies Ltd., Oxford, United Kingdom), following previously reported procedures (Colson et al., 2022).

### Metagenomic analyses of next-generation sequencing runs

Metagenomics results were obtained using the Kraken2 version 2.0.9 tool (Wood et al., 2014) against the database k2_standard as available on 2022-06-07.

### Monkeypox virus genome sequence analysis

Raw NGS reads were mapped against the MPX virus genome GenBank Accession no. ON563414.3 using minimap2 (Li, 2018). This 197,205 base pair-long genome was obtained from a patient sampled in Massachusetts, United States, in May 2022, and is used as a B.1 reference genome by the Nextclade tool v1.6.0 (https://nextstrain.org/monkeypox/hmpxv1) (Aksamentov et al., 2021). Different methods for variant calling were used according to the NGS technology. In case of Illumina reads, variant calling was performed with freebayes version 1.3.5 (Garrison and Marth, 2012) and in case of Nanopore reads it was performed with medaka 1.4.4, longshot v0.4.5 (https://github.com/pjedge/longshot) (Edge and Bansal, 2019), and artic_vcf_filter from the artic pipeline (https://artic.readthedocs.io/en/latest/). MPX virus consensus genomes were generated by putting a “N” at all positions covered by less than 4 reads for Illumina sequencing and less than 10 reads for Nanopore sequencing, and by considering all position tags as failed by artic_vcf_filter has “N”.

### Genome annotation

The set of MPX virus genes and proteins was predicted performed using the GeneMarkS tool (Besemer et al., 2001), with default parameters. Then, the set of amino acid sequences was used for a BLAST search (https://blast.ncbi.nlm.nih.gov/Blast.cgi) (Altschul et al., 1990) against the protein set from genome GenBank accession numbers NC_063383 and ON563414 and against the NCBI GenBank protein sequence database (nr) (Sayers et al., 2022); 0.01 was considered as significant threshold for the e-values.

### Phylogenetic analysis

The MPX virus genomes deposited in the GISAID sequence database (https://gisaid.org/) (Elbe et al., 2017) until 22^nd^ of August, 2022 and harboring the most mutations in common with those obtained in the present study were selected for their incorporation in the phylogenetic reconstruction within the limit of maximum two genomes (asides genomes obtained in the present study) by identified phylogenetic cluster. The genomes GenBank accession numbers NC_063383 and ON563414 were added to root the tree. Genome sequences were aligned with MAFFT v7.505 (Katoh et al., 2013). The genome region corresponding to nucleotide positions 150,551 to 150,576 (relatively to genome NC_063383) was excluded from the analysis because it contains a deletion and repeats that do no allow a robust assembly of NGS reads. The tree was built with the Iqtree tool v2.2.0.3 (Nguyen et al., 2015) with the -m GTR+R and -B 1000 options, then viewed with the MEGA software v11 (Kumar et al., 2018).

## ACKNOWLEDGMENTS

We are thankful to Marion Le Bideau, Claudia Andrieu and Priscilla Jardot for their technical help.

## DATA AVAILABILITY

Viral genomes obtained and analyzed in the present study have been deposited in the GenBank sequence database (https://www.ncbi.nlm.nih.gov/genbank/) (Sayers et al., 2022) under accession numbers OP382478-OP382495, and are available from the IHU Marseille Infection website (https://www.mediterranee-infection.com/19extst-ressources/donnees-pour-articles/genome-monkeypox/). They can also be retrieved online from the EpiPox sheet of GISAID sequence database (https://gisaid.org/) (Elbe et al., 2017) using the online search tool with “IHU” and “France” as keywords (GISAID identifiers are as follows: no. EPI_ISL_14863048, EPI_ISL_14863066, EPI_ISL_14863065, EPI_ISL_14863064, EPI_ISL_14863063, EPI_ISL_14863062, EPI_ISL_14863061, EPI_ISL_14863060, EPI_ISL_14863059, EPI_ISL_14863058, EPI_ISL_14863057, EPI_ISL_14863056, EPI_ISL_14863055, EPI_ISL_14863054, EPI_ISL_14863053, EPI_ISL_14863052, EPI_ISL_14863051, EPI_ISL_14863050, EPI_ISL_14621526, EPI_ISL_14621525, EPI_ISL_13308160, EPI_ISL_13308158, EPI_ISL_13052287).

## ETHICS

The present study has been approved by the ethics committee of University Hospital Institute (IHU) Méditerranée Infection (N°2022-040).

## AUTHOR CONTRIBUTIONS

P.C. and B.L.S. designed the study. P.C., G.P., J.D, C.B., N.W., S.B., J.CL., N.C., H.T.D., M.M., S.A., and B.L.S. provided materials, data or analysis tools. P.C., J.D., C.B., S.A., and B.L.S. analyzed the data. P.C., S.A., and B.L.S. wrote the first draft of the manuscript. All authors approved the final manuscript.

## SUPPLEMENTARY FIGURES

**Supplementary Figure S1.**
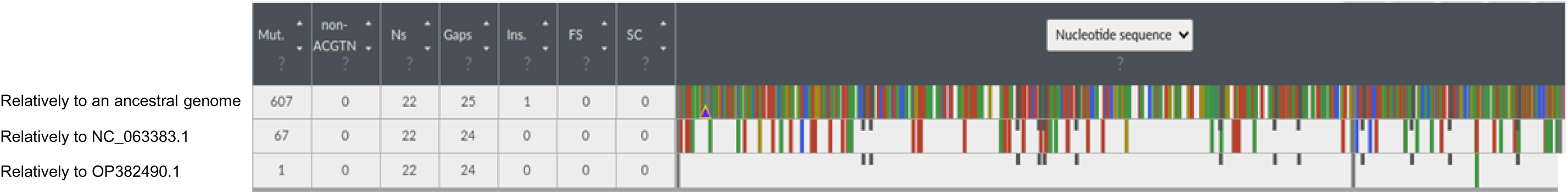
Screenshot of an output of the Nextclade tool dedicated to Monkeypox virus Nextclade (https://nextstrain.org/monkeypox/hmpxv1) (Aksamentov et al., 2021) showing the distribution of mutations along the Monkeypox virus genome hMPXV-IHU0001 (GenBank Accession no. OP382478.1) relatively to an ancestral genome and to reference genomes GenBank Accession no. NC_063383.1 and OP382490.1. The ancestral genome is a reconstructed sequence used by Nextclade tool dedicated to Monkeypox virus Nextclade (https://nextstrain.org/monkeypox/hmpxv1) (Aksamentov et al., 2021).

**Supplementary Figure S2.**
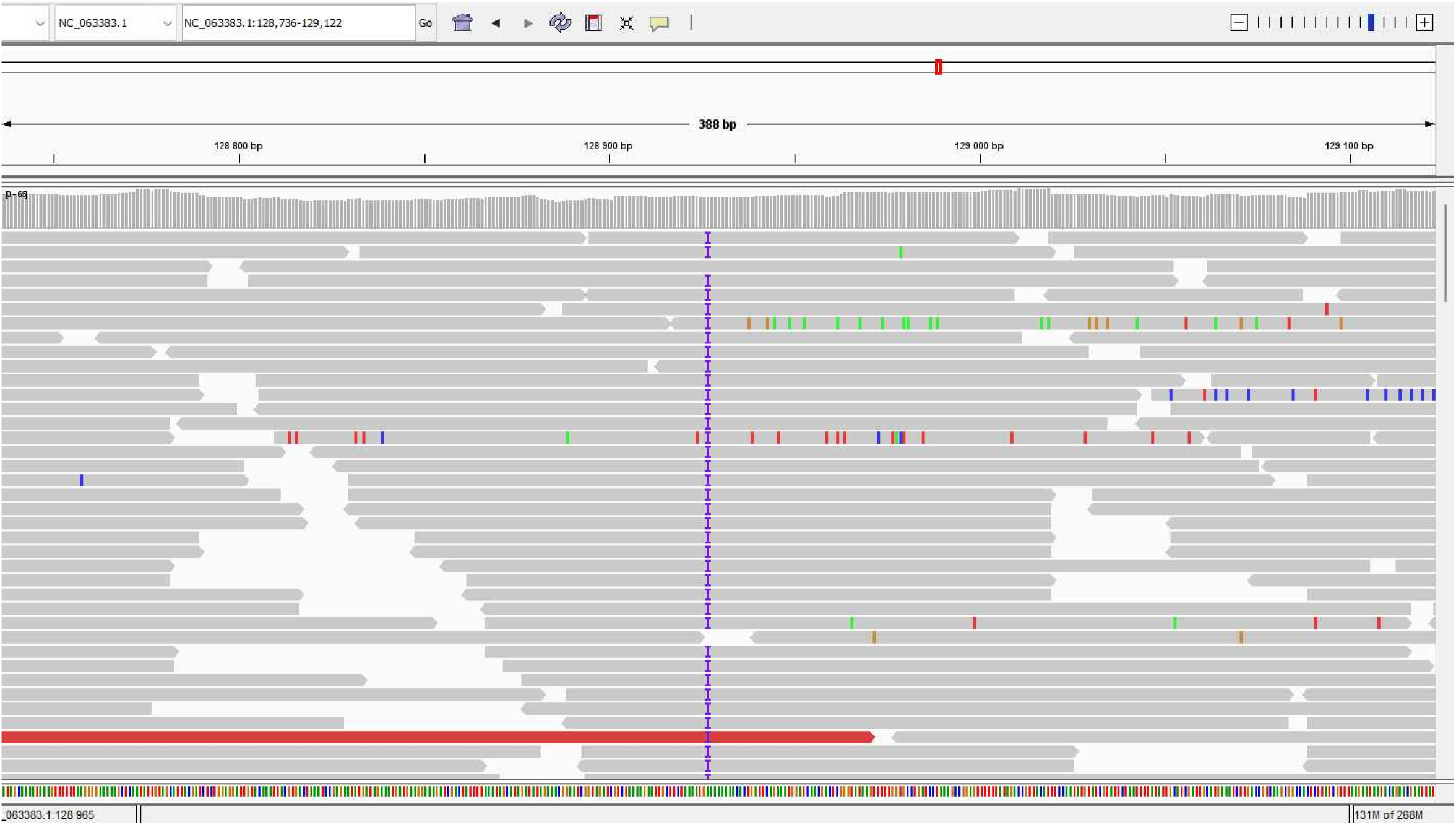
Screenshot of the assembly by mapping of next-generation sequencing reads for the Monkeypox virus genome hMPXV-IHU00013 (GenBank Accession no. OP382490.1) obtained in the present study, which shows the insertion of a nucleotide in the RNA polymerase subunit RPO132 encoding gene in 47 of 48 reads. Mapping was performed on the Monkeypox virus genome GenBank Accession no. NC_063383.1 using minimap2 (doi: 10.1093/bioinformatics/bty191).

**Supplementary Figure S3.**
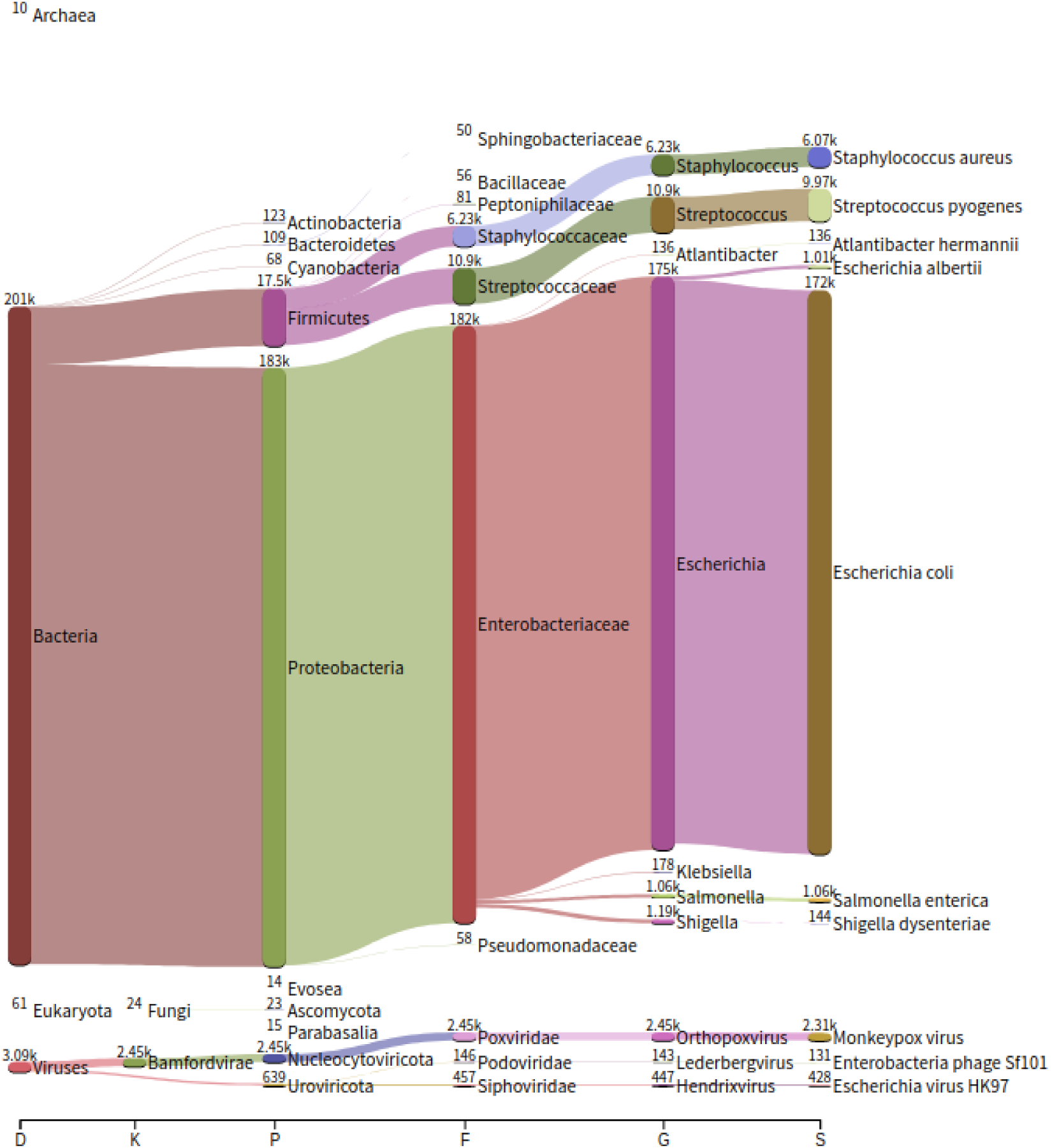
Screenshot of a Kraken2 output for metagenomic analysis of next-generation sequencing reads obtained for the penis lesion from which Monkeypox virus genome hMPXV-IHU0001 (GenBank Accession no. OP382478.1) was obtained in the present study, which shows the presence of 6,070 and 9,970 reads identified as most similar to genomes from *Staphylococcus aureus* and *Streptococcus pyogenes*, respectively.

## SUPPLEMENTARY TABLE

**Supplementary Table S1.**
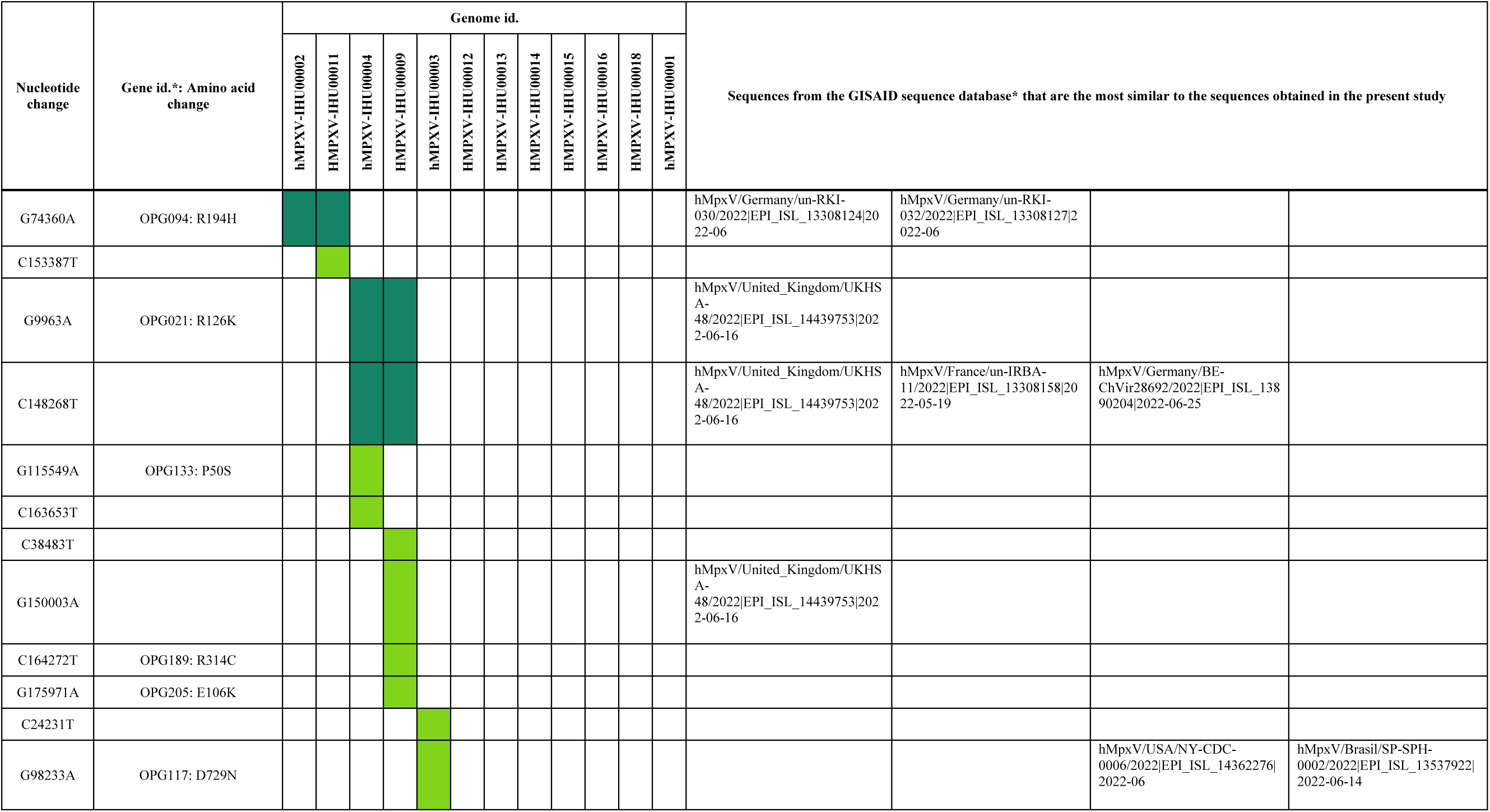
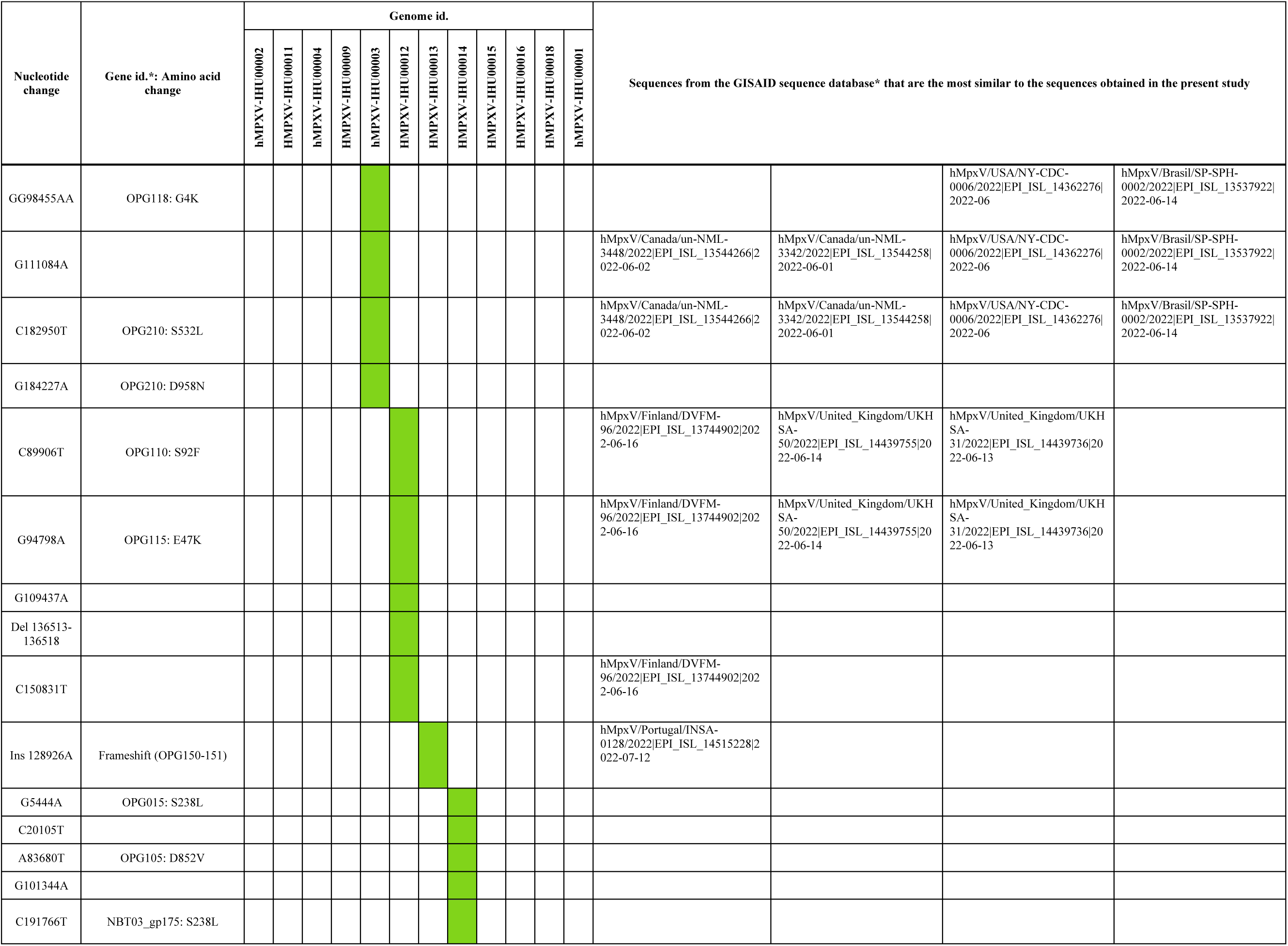
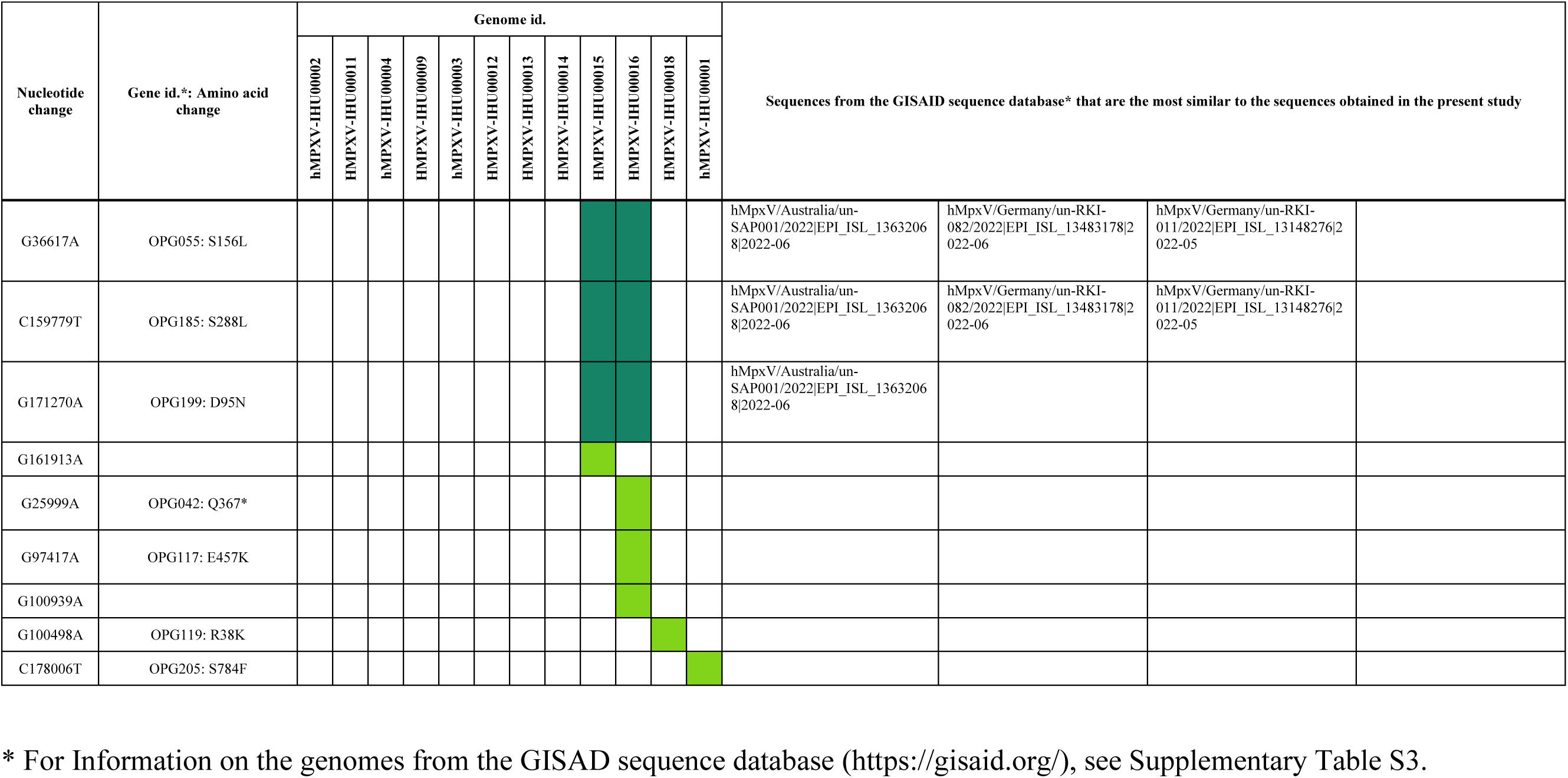
Nucleotide and amino acid change in Monkeypox virus genomes obtained in the present study relatively to genome GenBank Accession no. ON563414.3, for the genomes that harbor at least one mutation, and genomes the most similar in the GISAID sequence database (https://gisaid.org/)

**Supplementary Table S2.**
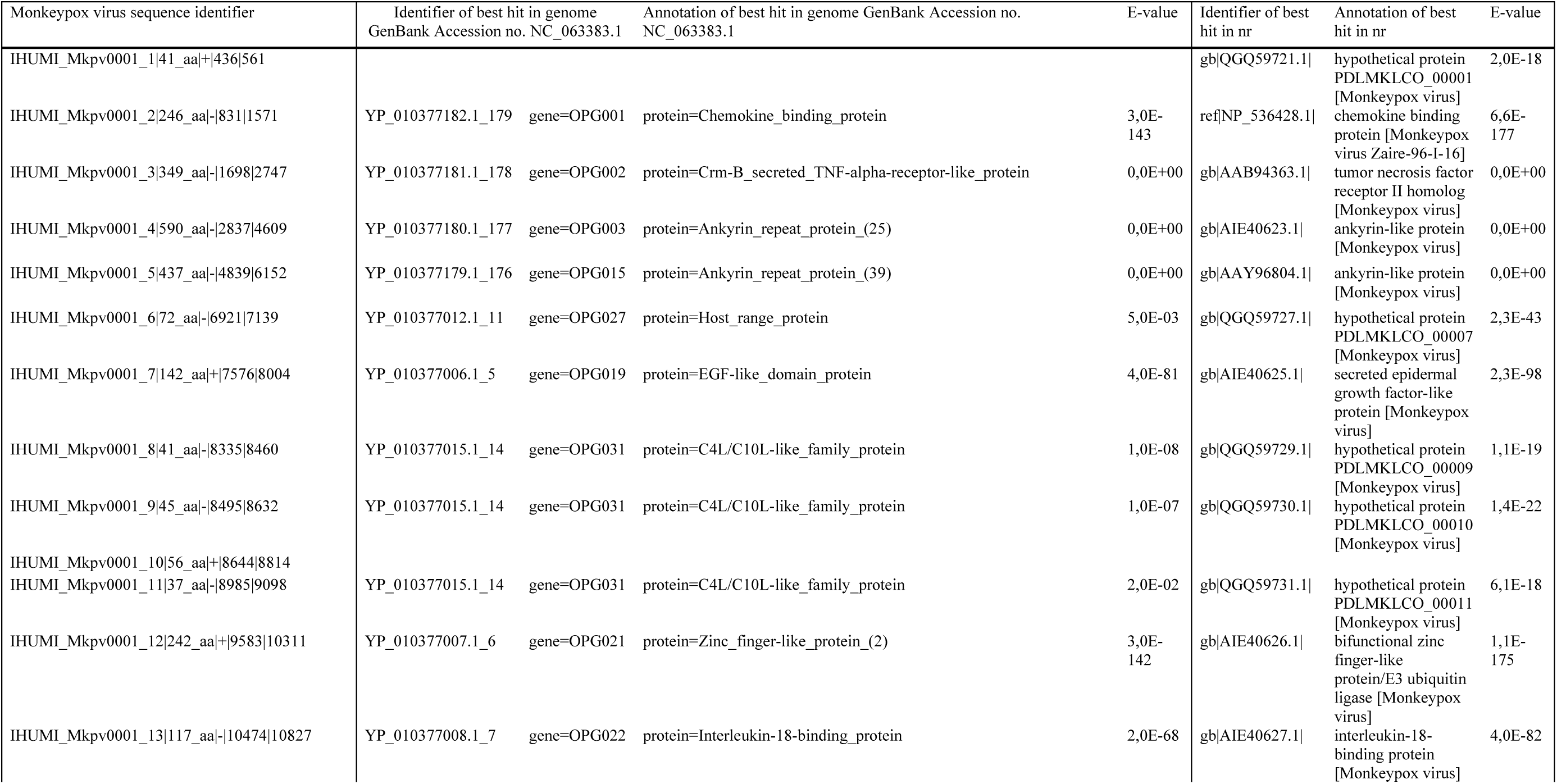
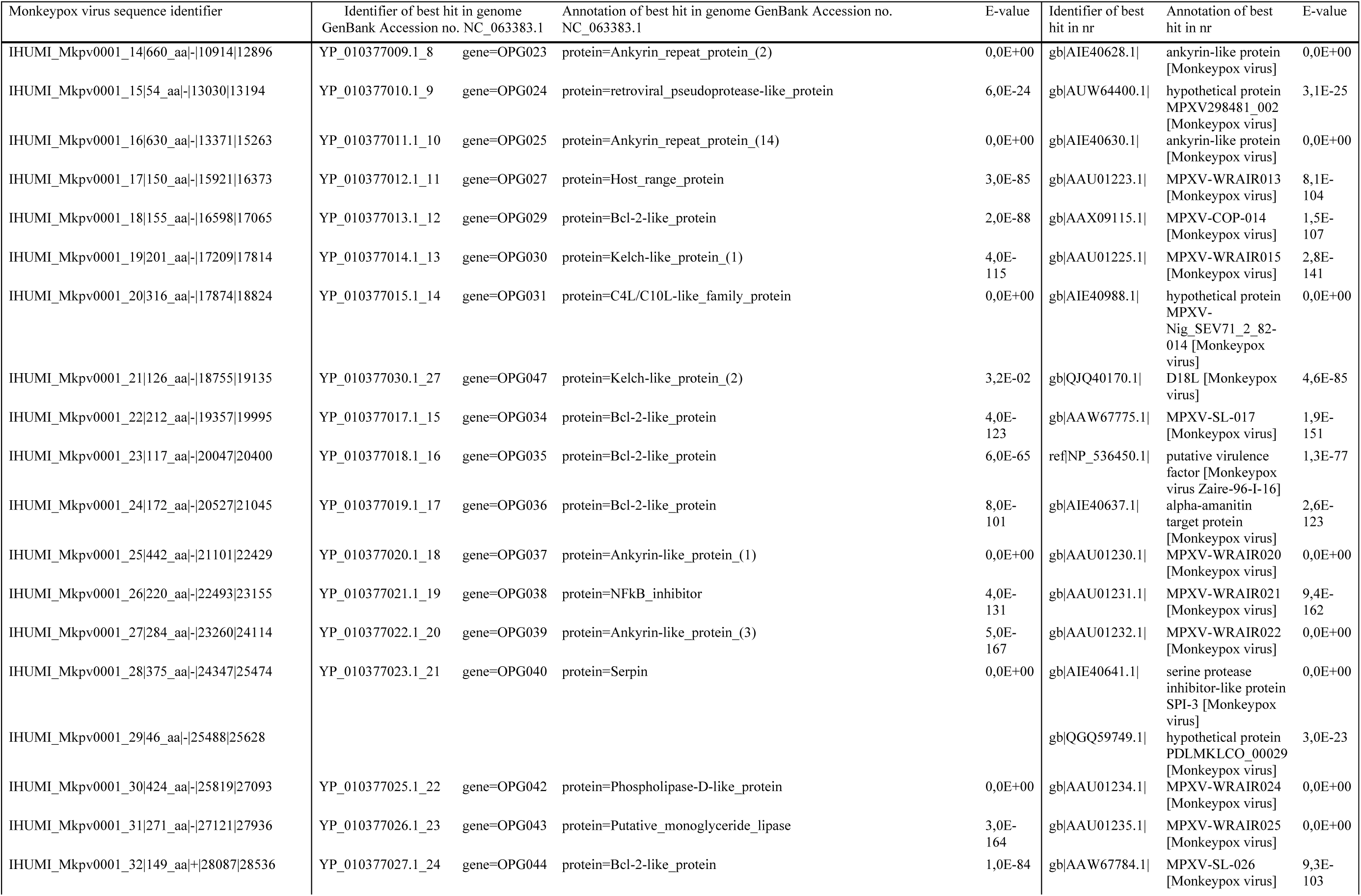
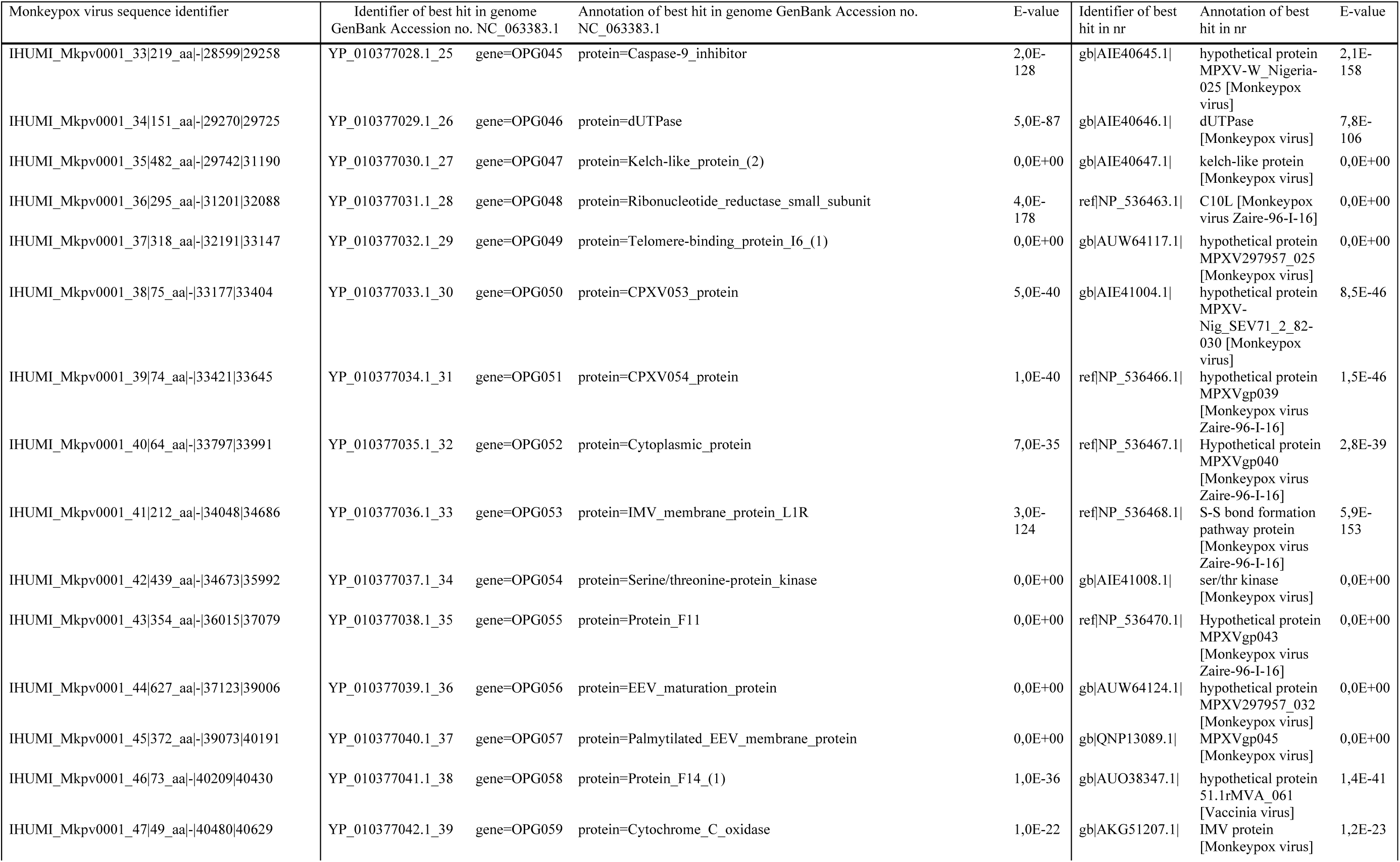
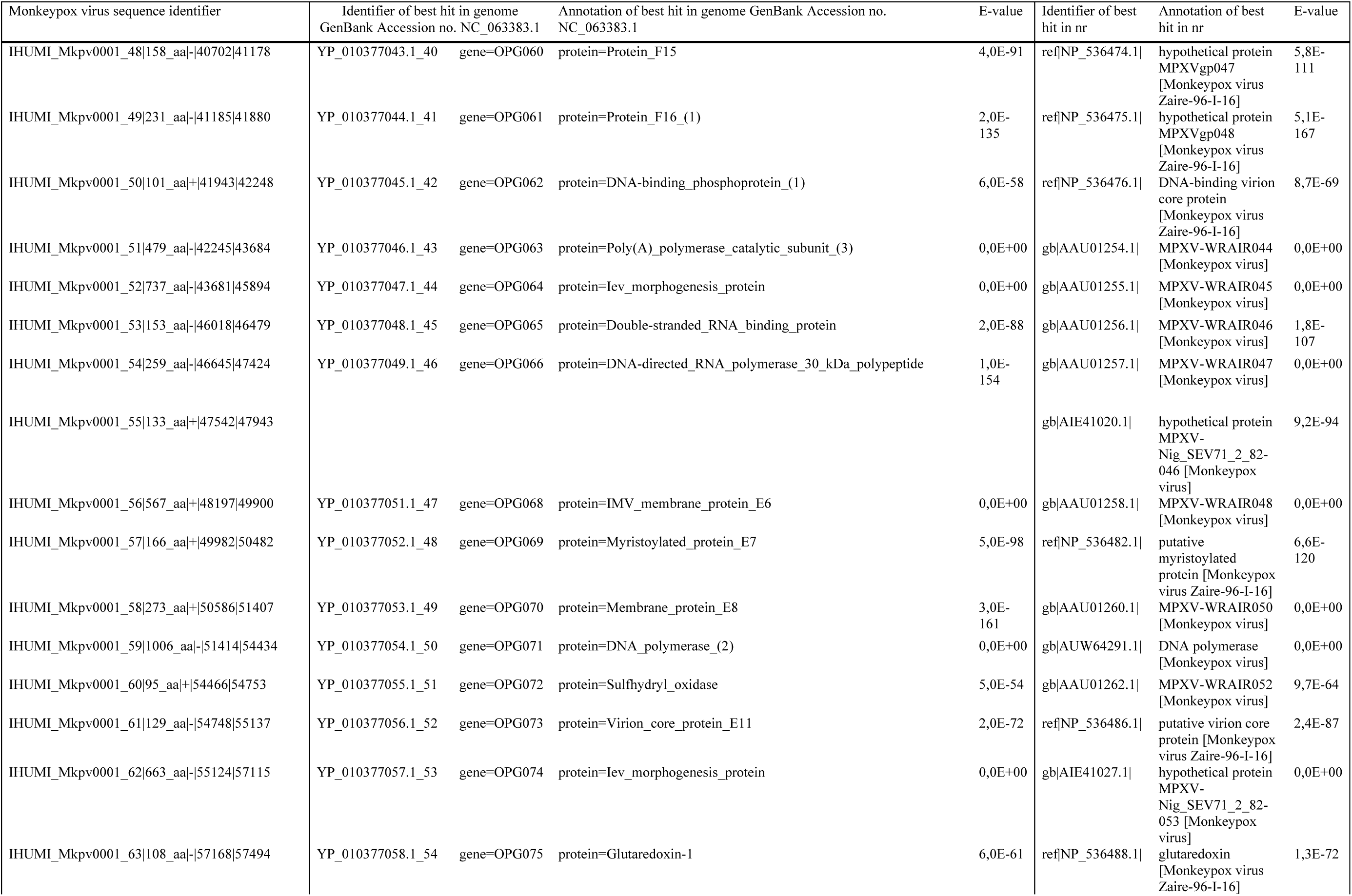
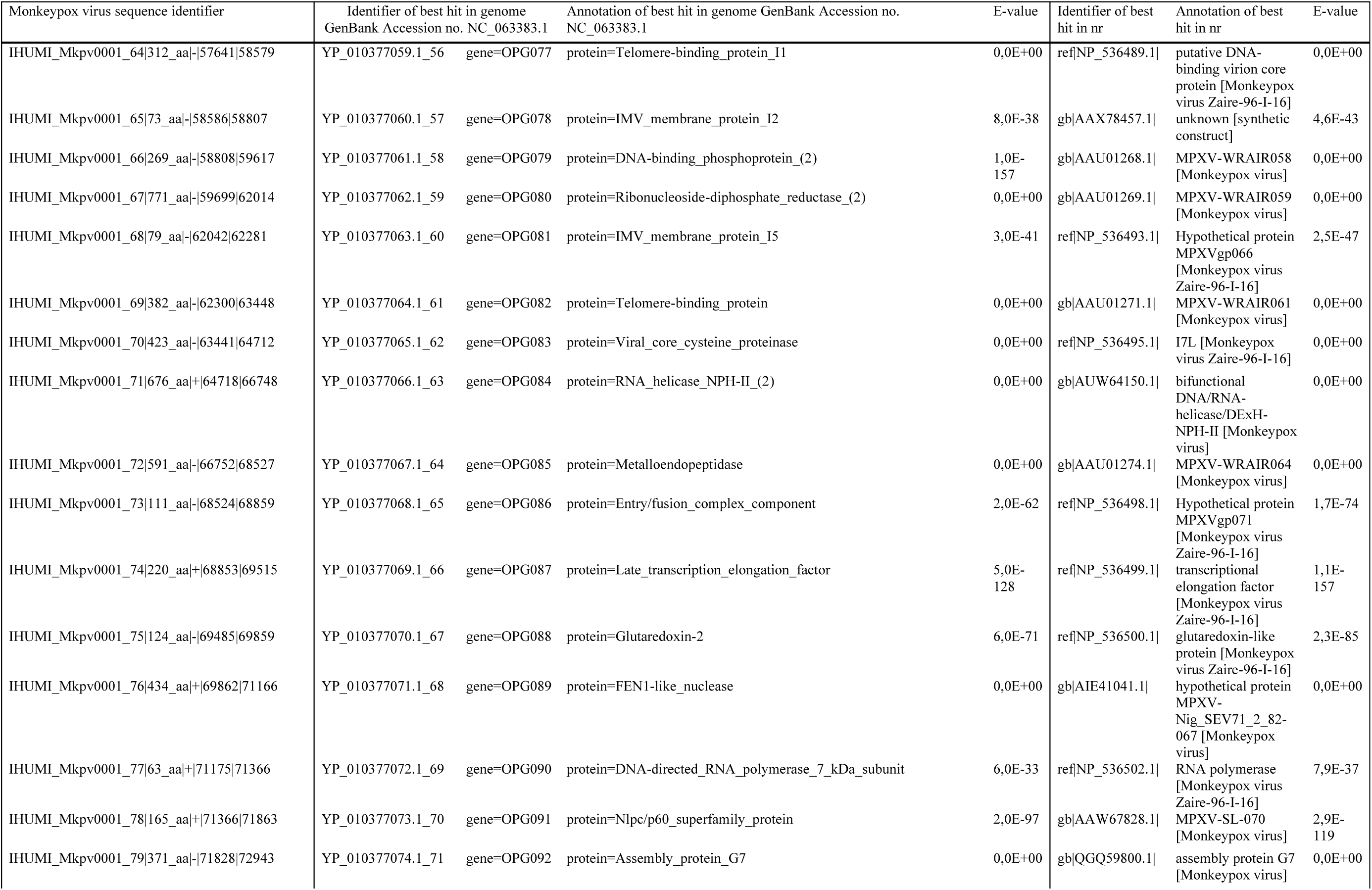
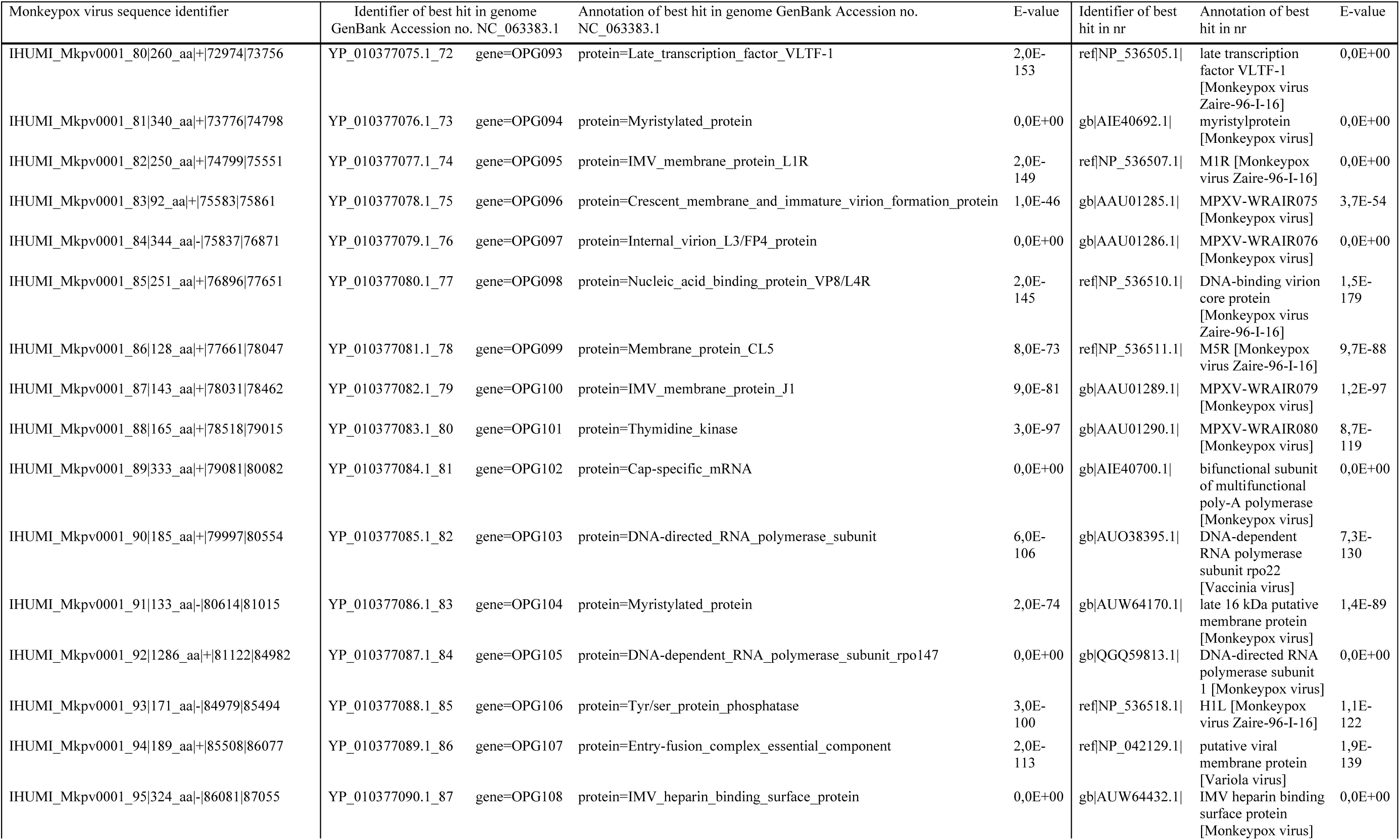
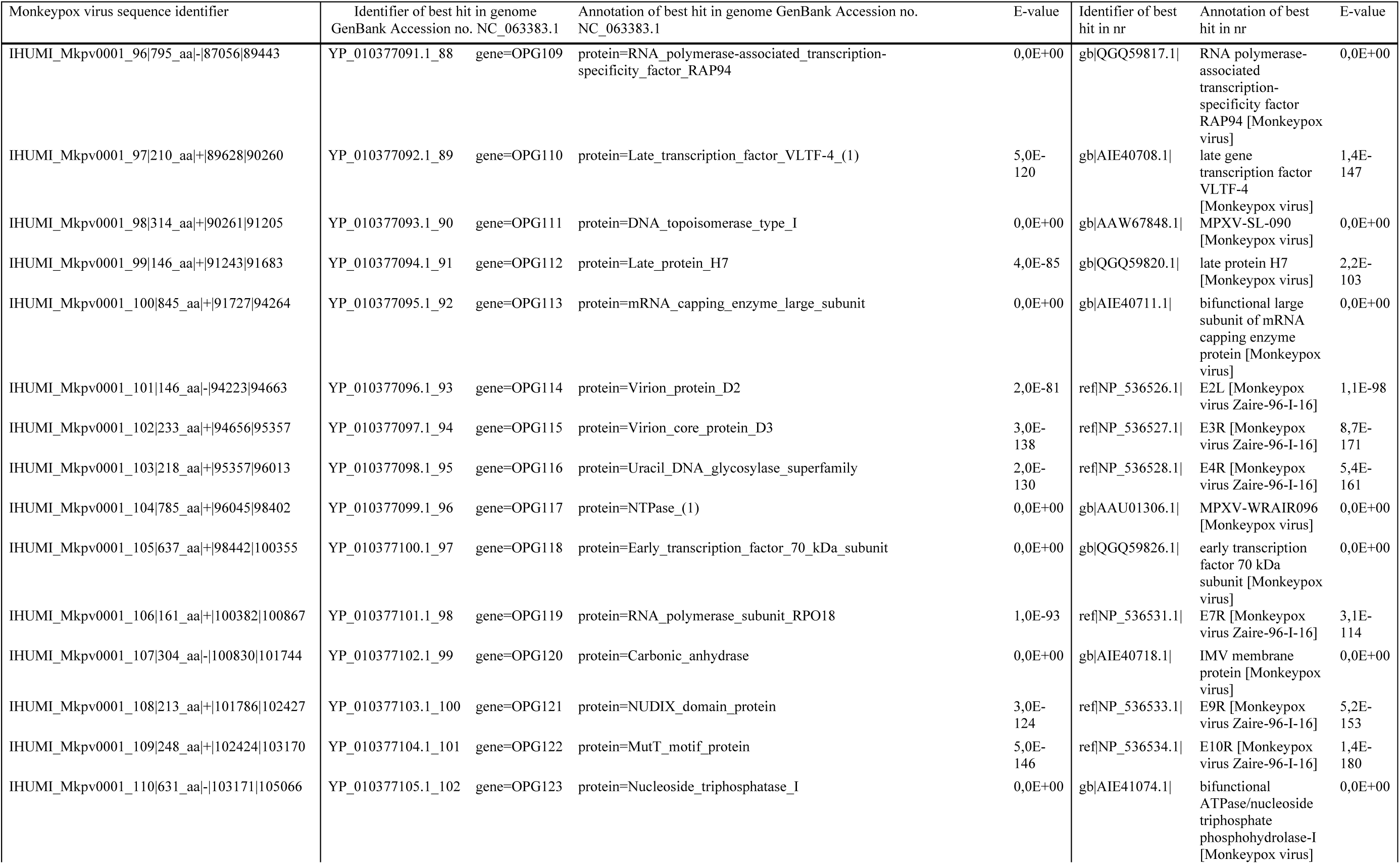
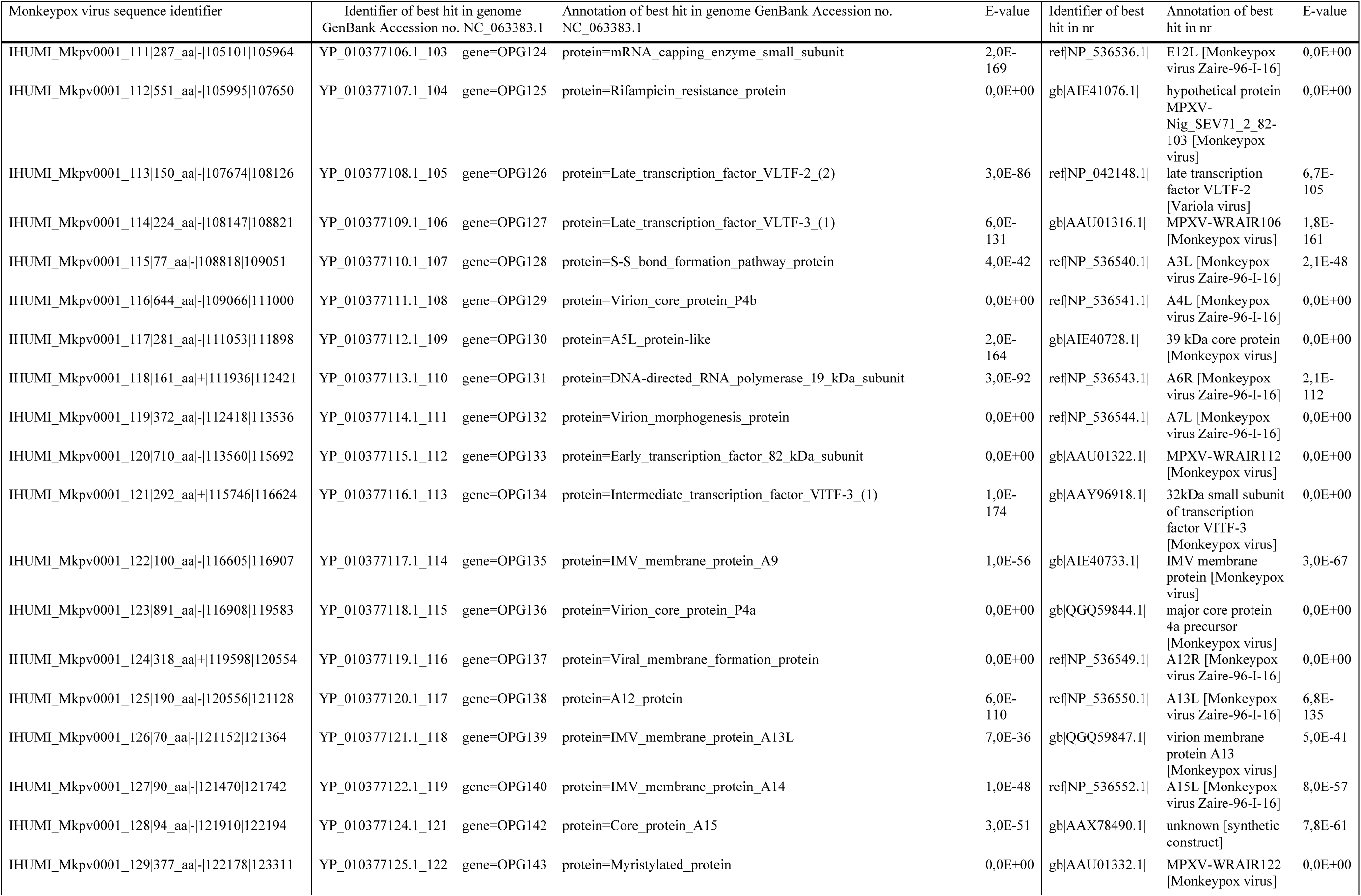
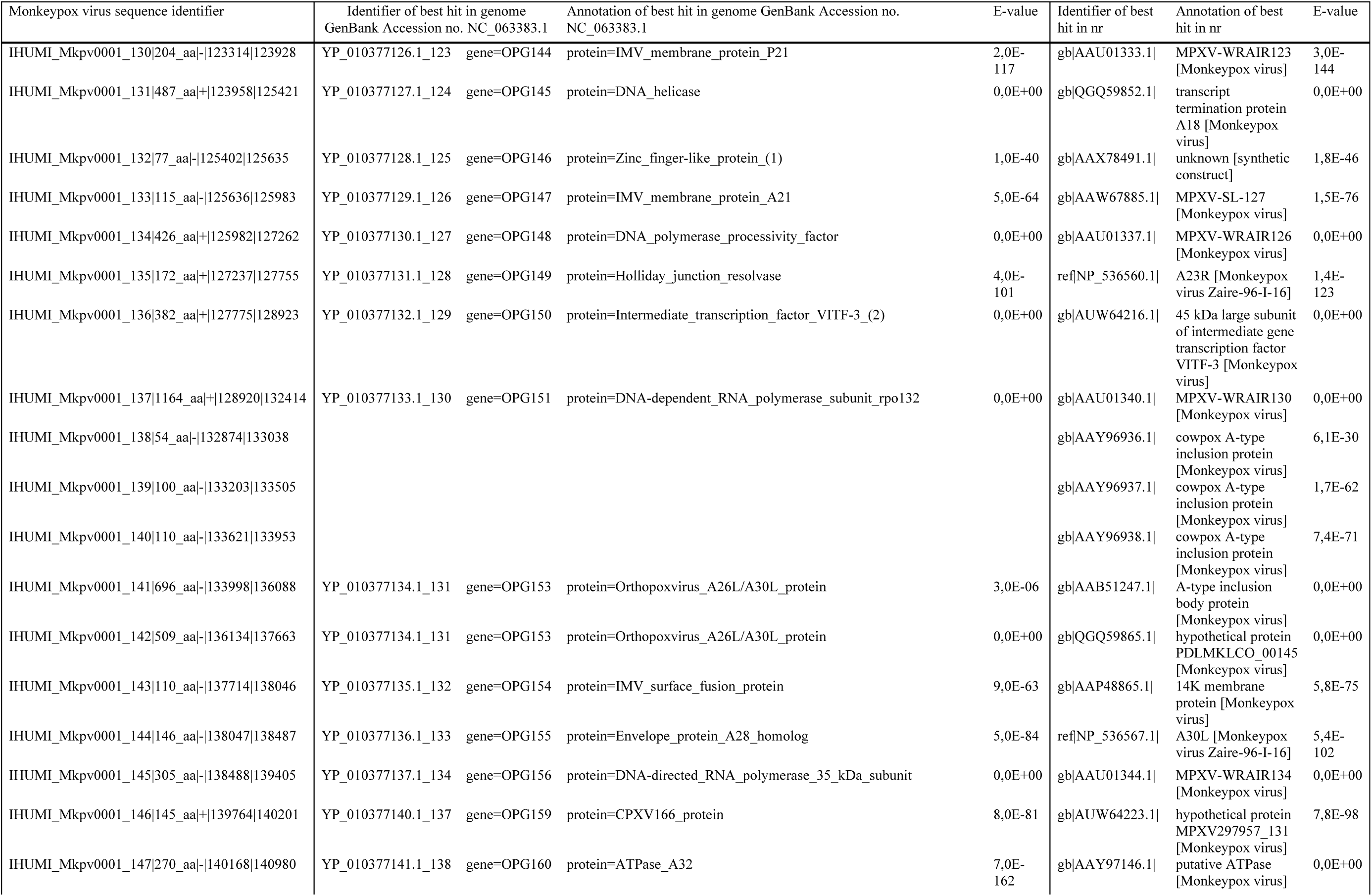
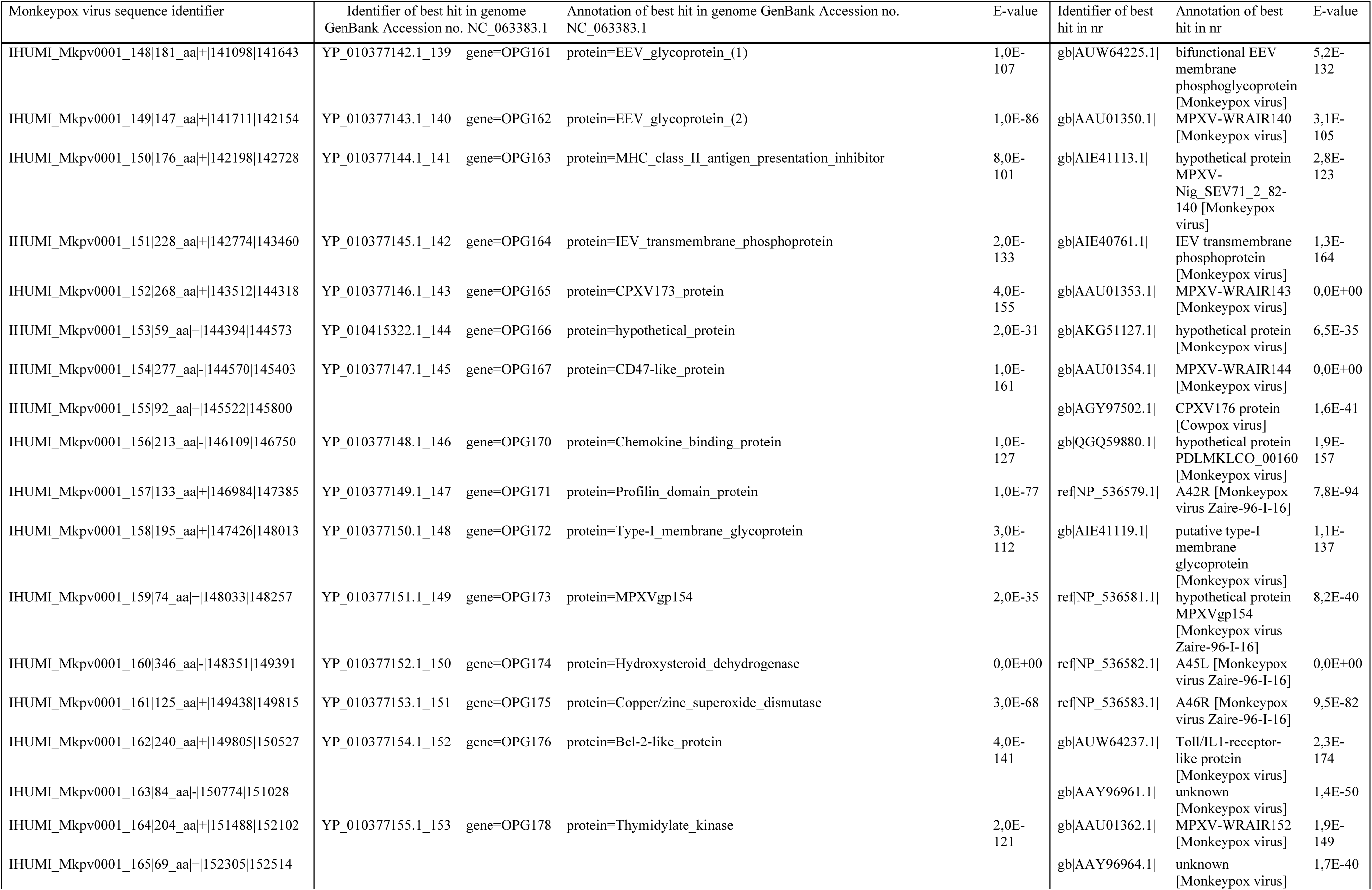
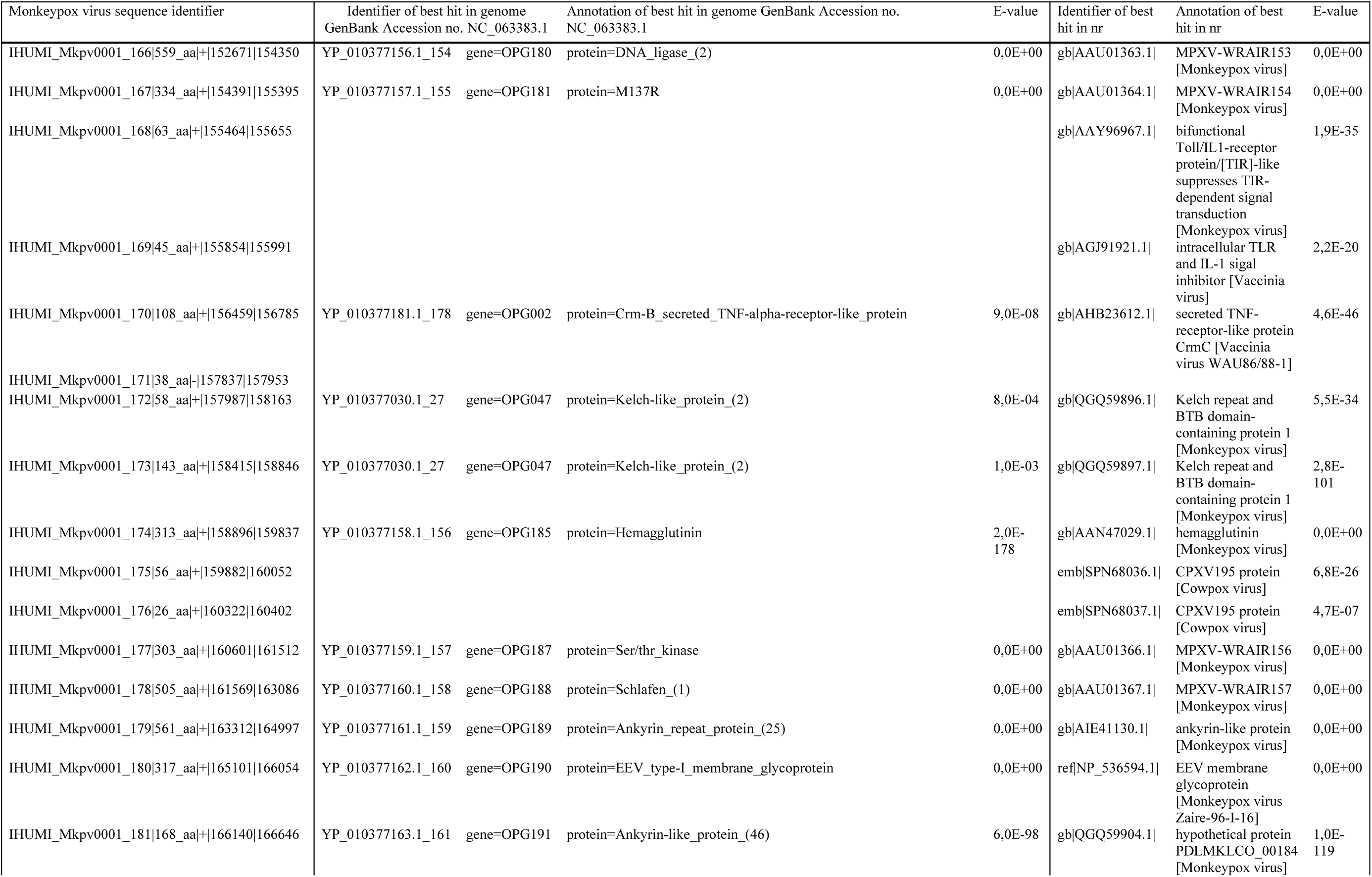
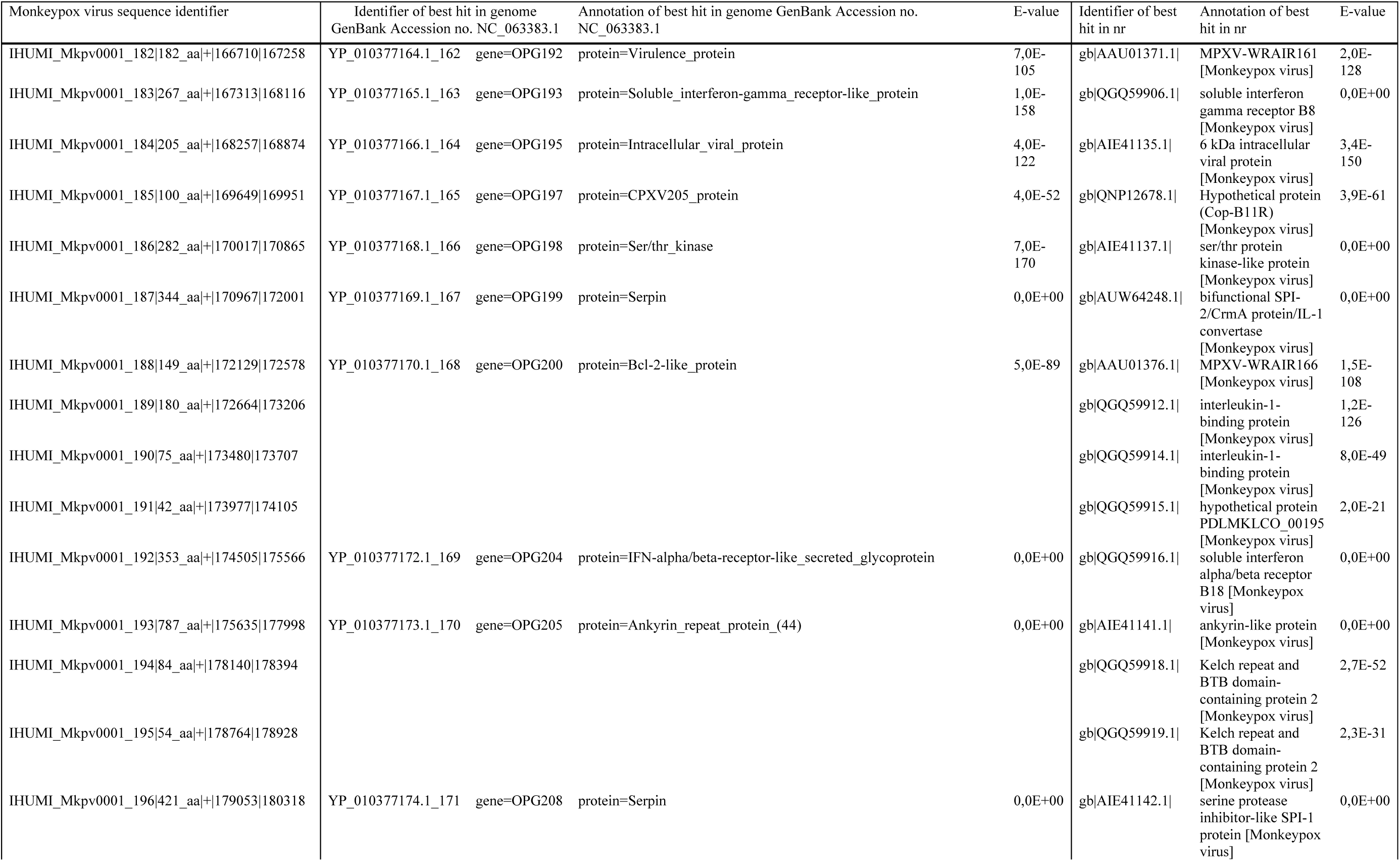
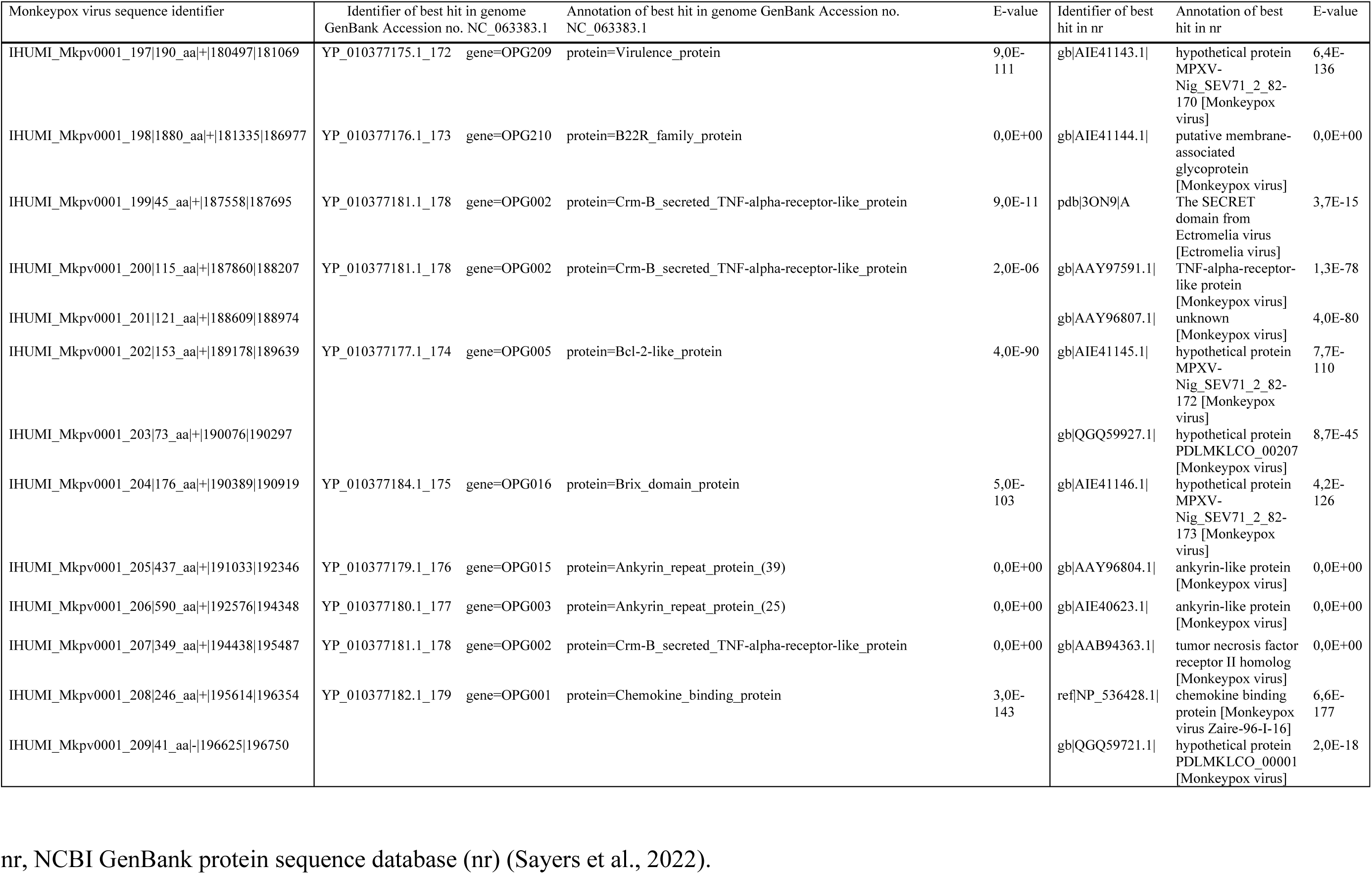
Annotation of the Monkeypox virus genome (hMPXV-IHU00001; GenBank Accession no. OP382478.1; 208 gene products) that we obtained using GeneMarkS (Besemer et al., 2001)

**Supplementary Table S3.**
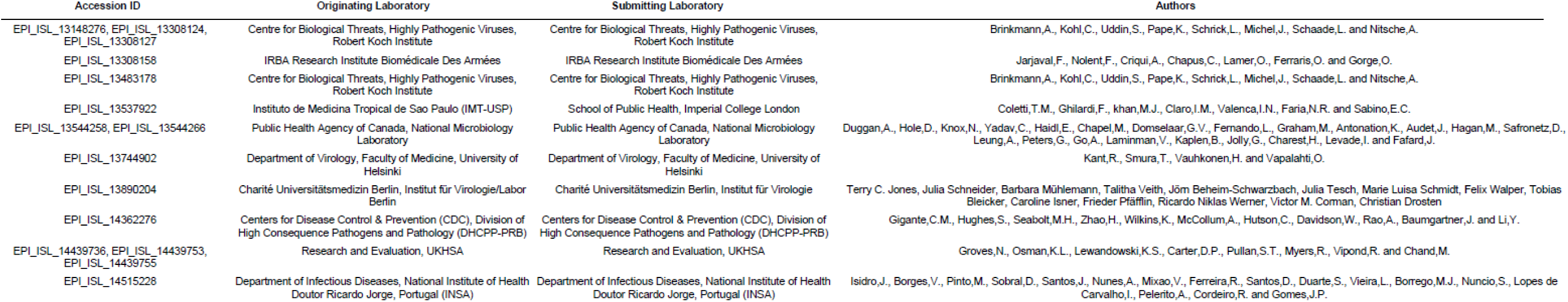
Acknowledgment table for genome sequences from the GISAID database quoted in the present study (https://gisaid.org/) (Elbe et al., 2017)

